# Conflicting Timelines: Exploring patterns of mito-nuclear discordance in divergence estimates among tetrapods

**DOI:** 10.1101/2024.12.02.626515

**Authors:** Ekadh Ranganathan, Praveen Karanth

## Abstract

Phylogenetic studies across a range of tetrapod groups have historically utilised either mitochondrial DNA, concatenated mito-nuclear matrices, or nuclear loci to infer divergence estimates. As such, the discordance between estimates inferred using differing data types has been a topic of interest over the past decade. Although several studies have looked into divergence disparities in smaller taxonomic groups, often with mixed results, a systematic investigation of the pattern of mito-nuclear divergence discordance in tetrapods across deep-time remains to be undertaken. In this study, we aimed to quantify the extent of divergence disparity inferred by the aforementioned data types in each of the four major tetrapod groups, namely primates (Order *Primates*), birds (Class *Aves*), squamates (Order *Squamata*), and anurans (Order *Anura*), while controlling for calibration strategies, taxon sampling and other *a priori* distributions. We also calculated substitution saturation for all groups across data types in order to elucidate its role in generating unreliable divergence estimates. Our findings indicate that mitochondrial estimates consistently underestimate basal divergence times and overestimate recent divergences across all groups (apart from anurans) as compared to the nuclear datasets. We also find that divergence times estimated using concatenated matrices skew in favour of the nuclear tree. Furthermore, substitution saturation is substantial in all of the mitochondrial datasets across groups, and interestingly present in the nuclear dataset for Anurans, resulting in a reduction in overall mito-nuclear divergence disparity for the crown ages of the group. These results call for a revisit of divergence dates estimated using mitochondrial DNA, while advocating for saturation testing prior to divergence dating and highlighting the inherent bias in divergence dates estimated using concatenated mito-nuclear matrices.

---

The discovery and implementation of molecular clocks in phylogenetics has allowed for the estimation of divergence dates across a range of taxonomic groups. In tetrapod groups, these divergence dating studies have historically employed either mitochondrial DNA markers (mtDNA), nuclear DNA markers (nDNA), or a concatenated mix of the two marker types (Huelsenbeck et al., 1996; Hwang and Kim, 1999). However, the interchangeable usage of genetic markers from this pool of competing data types has yielded vastly incongruent estimates of divergence across the phylogenetic tree (Mulcahy et al., 2012; Toews and Brelsford, 2012; Smith and Klicka, 2013; Bernardo et al., 2019).

Mitochondrial markers have been the primary choice for phylogenetic analyses predominantly due to their short coalescent time, high variation, lack of recombination and relative ease of isolation and sequencing (Rubinoff and Holland, 2005; Zheng et al., 2011). However, questions have been raised regarding their utility as markers for molecular dating due to their rapid substitution rates, particularly at 3rd codon positions, which can often bias divergence estimates due to the adverse effects of substitution saturation (Gojobori, 1983; Blouin et al., 1998; Xia, 1998; Nilsson et al., 2010). On the other hand, nuclear markers (especially exonic protein-coding nuclear markers) accumulate fewer substitutions through time, and are thus broadly unaffected by saturation to the same extent as mtDNA (Brower and DeSalle, 1998; Irestedt et al., 2001; Springer et al., 2001).

## Mixed effects of saturation on divergence estimates

Substitution saturation is the process by which multiple substitutions accumulate at singular nucleotide sites in an alignment (Lopez et al., 1999; Philippe and Forterre, 1999; Xia et al., 2009). These rapid substitutions occur such that the observed genetic distances across the alignment can no longer reflect historical variation, even when inferred by a complex substitution model like GTR (Arbogast et al., 2002). Thus, the phylogenetic information at these sites is lost, as the number of substitutions are generally underestimated in cases of severe saturation (Xia et al., 2003). As such, when considering deep-time divergences wherein sequences tend to experience significant saturation effects, this loss of phylogenetic information can thus cause an underestimation of true divergence times among ancestral taxa (Wilke et al., 2009), as exemplified in a number of studies (Phillips and Penny, 2003; Brown et al., 2008a; Ho and Phillips, 2009; Near et al., 2012; Bocak et al., 2014).

Furthermore, Arbogast et al. (2002) and Zheng et al. (2011) argued that divergence estimates ascertained from data that has undergone saturation would skew towards fossil calibration points (due to the loss of phylogenetic signal), such that dates younger than a singular calibration point will be overestimated, and dates older than the calibration point will be underestimated. However, Wilke et al. (2009) argued that divergences among closely related taxa are plagued by factors other than substitution saturation such as ancestral polymorphisms and power gaps, which are more to blame for the inaccurate estimation of shallower nodes. Consequently, the pattern of discordance caused by differential saturation across data types and especially timescales still remains poorly explored for most taxonomic groups, including most tetrapods.

Although the underestimation of divergence estimates from saturated data has been theoretically proven, only a handful of studies have actually shown that dates estimated using mtDNA markers underestimate true divergence times, namely in birds and caenophidian snakes (Brown et al., 2008b; Lukoschek et al., 2011). Other studies have discovered an inverse pattern of mtDNA overestimation across the tree, like in salamanders, squamates, and rhesus macaques (Melnick et al., 1993; Ho et al., 2005; Brandley et al., 2010; Zheng et al., 2011; Mulcahy et al., 2012). These conflicting outcomes could be explained by a plethora of *a priori* differences in parameter choices, especially differing calibration strategies. Nonetheless, on account of such uncertainties between theoretical and empirical tests of divergence discordances, it would be prudent to assemble dated phylogenies for a range of tetrapod groups while controlling for confounding factors such as fossil calibrations and taxon sampling, in order to understand the true extent of divergence discordance between data that is likely saturated (mtDNA), and data that generally does not undergo extensive saturation through time (nDNA), and whether the pattern holds true across groups.

## Divergence dating using concatenated matrices

Although the prevalence of studies utilising purely mitogenomes to infer deep-time divergence estimates has declined in the past decade with the advent and increased availability of High-Throughput Sequencing (Delsuc et al., 2005; Patané et al., 2018; Braun et al., 2019; Steenwyk et al., 2023), concatenated dating using mtDNA and nDNA markers is still a widespread practice (Jønsson et al., 2016; McCraney et al., 2020; Henríquez-Piskulich et al., 2024). These concatenated dating studies have helped provide insight into evolutionary processes in a range of taxonomic groups for which genetic data was otherwise scarce, such as for rodents, bats and bees (Fabre et al., 2012; Bartlett et al., 2013; Cardinal and Danforth, 2013). However, there is a dearth of research addressing issues caused by the potential skew of divergence estimates towards mtDNA or nDNA in such alignments. Fisher-Reid and Wiens (2011) looked at skewness in concatenated alignments in terms of its effects on species relationships and branch lengths. However, *a priori* parameters like taxon sampling were not standardised across the 13 groups studied, and the study did not reflect the effects of concatenation on divergence times. Thus, this study also aimed to understand the skew of concatenated divergence estimates (rather than species relationships) towards either purely mtDNA or nDNA estimates across tetrapod groups while employing a similar coverage of both marker types in each alignment.

This study thus has three primary objectives: (1) to understand the pattern and extent of divergence discordance between mtDNA and nDNA datasets across large tetrapod groups, (2) to investigate the role of substitution saturation in explaining the pattern of discordance seen across data types, and (3) to explore the extent of skew of divergence estimates of mito-nuclear matrices towards mtDNA or nDNA estimates.

## Materials and Methods

### Taxon Sampling and Alignment composition

Secondary alignments were sourced from four widely-studied tetrapod groups for which mtDNA and nDNA sequence data were readily available (refer to Table 1), namely the Order *Primates*, Class *Aves*, Order *Squamata* and Order *Anura*. Since clades with vastly differing crown ages and number of taxa cannot be directly compared in terms of divergence discordance, estimates from the phylogenies of Order Primates and Class Aves (crown ages: 70-100 Ma) were compared independently of Order Squamata and Order Anura (crown ages: 170-200 Ma). Crown groups of different ages were chosen to better explore discordance at different timescales. Within each group, the taxa used for divergence dating remained consistent across the mtDNA, nDNA and concatenated alignments. Non-overlapping species between nDNA and mtDNA datasets were also included as long as they belonged to the same genus (see Supplementary File S1). The aforementioned non-overlapping species were directly concatenated when creating the mito-nuclear matrices. Lastly, in almost all groups, the mtDNA and nDNA datasets already consisted of overlapping outgroup taxa, which were left unchanged for this study (see Table 1, Supplementary File S1 for detailed information on taxon sampling). However, the mtDNA dataset for birds lacked an outgroup entirely, and thus an *Alligator mississippiensis* mitochondrion sequence was sourced from Genbank (Accession no.: AF069428.1) to ensure that the overlap in outgroup taxa between nDNA and mtDNA alignments was maintained across all groups.

**Table 1.** Alignment details for each group and data type.

| Group | Data type | Number of taxa | Alignment length(bp) | Outgroups | Variable sites in matrix(bp) | Dataset |
| --- | --- | --- | --- | --- | --- | --- |
| Order *Primates* | mtDNA | 54 | 13756 | *Galeopterus variegatus* | 8592 | Pozzi et al. (2014) |
|  | nDNA |  | 27436 |  | 8580 | Perelman et al. (2011) |
| Order *Squamata* | mtDNA | 139 | 8616 | *Sphenodon punctatus* | 6064 | Pyron et al. (2013) |
|  | nDNA |  | 33717 |  | 6069 | Wiens et al. (2012) |
| Order *Anura* | mtDNA | 119 | 13329 | *Andrias davidianus*,<br>*Batrachuperus sp.*,<br>*Gallus gallus*,<br>*Ichthyophis bannanicus* | 5890 | Portik et al. (2023) |
|  | nDNA |  | 88302 |  | 5895 | Feng et al. (2017) |
| Class *Aves* | mtDNA | 82 | 12947 | *Alligator mississippiensis* | 7938 | Arcones et al. (2021) |
|  | nDNA |  | 52383 |  | 7968 | Hackett et al. (2008) |

Alignments sourced from existing datasets were re-aligned and pruned where necessary in MEGA 11 (Tamura et al., 2021), and subsequently exported for downstream analyses. The partitioning scheme utilised for divergence dating across mtDNA and nDNA alignments was codon-specific, with all 1st, 2nd, and 3rd codon positions from exonic protein-coding regions forming three independent partitions. Intronic segments in the primate and bird nuclear datasets formed a single separate partition. Initial analyses for all groups and data types utilised a gene and codon-specific partitioning scheme, which was later discarded due to difficulties reaching stationarity across MCMC runs and poor effective sample sizes (ESS) across parameters, likely caused by overparameterisation.

Furthermore, by employing a simple codon-specific scheme in all analyses, variance in the number of partitions across groups could also be controlled for. As such, all mtDNA and nDNA analyses for squamates and anurans consisted of three partitions, while all primates and birds analyses consisted of four partitions (including intronic segments).

Mito-nuclear matrix alignments were assembled by directly concatenating the respective nDNA and mtDNA alignments for each group. However, since the nDNA datasets consisted of a substantially larger number of genetic markers (and thus informative sites) which could introduce prior bias into the skew of divergence estimates, the nDNA datasets were pruned to include a smaller random subset of markers such that the number of variable sites between mtDNA and nDNA alignments were approximately the same (see Table 1). Furthermore, each respective alignment was checked for missing data percentages (not shown) to ensure that all mtDNA and nDNA markers in a concatenated matrix possessed similar data coverage. All final dated phylogenies can be found in Supplementary File S3.

### Divergence Dating

All divergence dating runs were carried out in BEAST v2.7 (Bouckaert et al., 2019) on CIPRES (Miller et al., 2011). General time-reversible (GTR) substitution models (Tavaré and Miura, 1986) with four discrete gamma rate categories were chosen for all partitions. Most of the original datasets sourced for this study were dated using the GTR model in their respective studies, and it remains the most complex model of sequence evolution, thus facilitating our choice of the model for subsequent molecular dating. Fossil calibrations for each group were also chosen from either the mtDNA or nDNA study from which the data was sourced, depending on the strength and justification for each calibration point. Details regarding fossil calibration choices for this study are available in Supplementary File S1. Log-normal prior distributions were applied for each calibration point, as they remain the most suitable and widely used distributions for most fossil calibrations (Ho and Phillips, 2009). All calibrations were kept identical across nDNA, mtDNA and concatenated runs. The Yule model was employed with a relaxed log-normal clock prior to allow for rate heterogeneity across evolutionary time. A minimum of 3 MCMC replicates were carried out for each dataset for 400,000,000 generations each, as individual runs were analysed in Tracer v1.7.2 (Rambaut et al., 2018) to ensure ESS values above 200 for all parameters. After assessing chain convergence and ensuring adequate mixing across runs, log and tree files were combined using LogCombiner. Divergence estimates for each node in mtDNA, nDNA and concatenated runs were tabulated and plotted against each other using the ape and ggplot packages in R (R Core Team et al., 2013; Wickham, 2016; Paradis and Schliep, 2019). Nodes that were dissimilar across phylogenies of the differing data types were manually pruned to incorporate exclusively common nodes across runs to make for a fair comparison.

### Substitution Saturation Analysis

The extent of saturation in each dataset was determined by plotting observed against expected genetic distances as per Philippe et al. (2004). TN93 distance matrices (Tamura and Nei, 1993) and uncorrected p-distance matrices were assembled in MEGA11. Plots were made in R for individual codon positions in each dataset (following the partitioning scheme used for divergence dating in BEAST), with linear regression models fitted to identify differences in saturation slopes between nDNA and mtDNA datasets.

Since the extent of saturation is greatest in 3rd codon position plots across all groups and datasets, only 3rd codon saturation plots are displayed (all plots including 1st and 2nd codon position comparisons are available in Supplementary File S2).

## Results

### Divergence discordance

Primate divergence estimates for the mtDNA vs. nDNA, mtDNA vs. concatenated and concatenated vs. nDNA datasets are shown in Figure 1. The solid red line with a slope of one demarcates congruent dates between the data types. As such, divergence estimates that fall below the line indicate older mitochondrial estimates, and vice versa for estimates that fall above the line. When directly comparing mtDNA and nDNA estimates, the mtDNA dates tended to overestimate ages for shallow nodes, while most basal divergences greater than 20 Ma tended to favour a significantly older nuclear estimate (Fig 1a). A similar pattern was observed for mtDNA vs. concatenated divergence estimates, with mtDNA overestimating ages of younger nodes while underestimating the ages of older nodes compared to the concatenated dataset (Fig 1b). Interestingly, the concatenated mito-nuclear matrix dates showed a strong correlation with nDNA estimates despite controlling for the number of variable sites in both alignments prior to concatenation (Fig 1c).

**Fig. 1.**
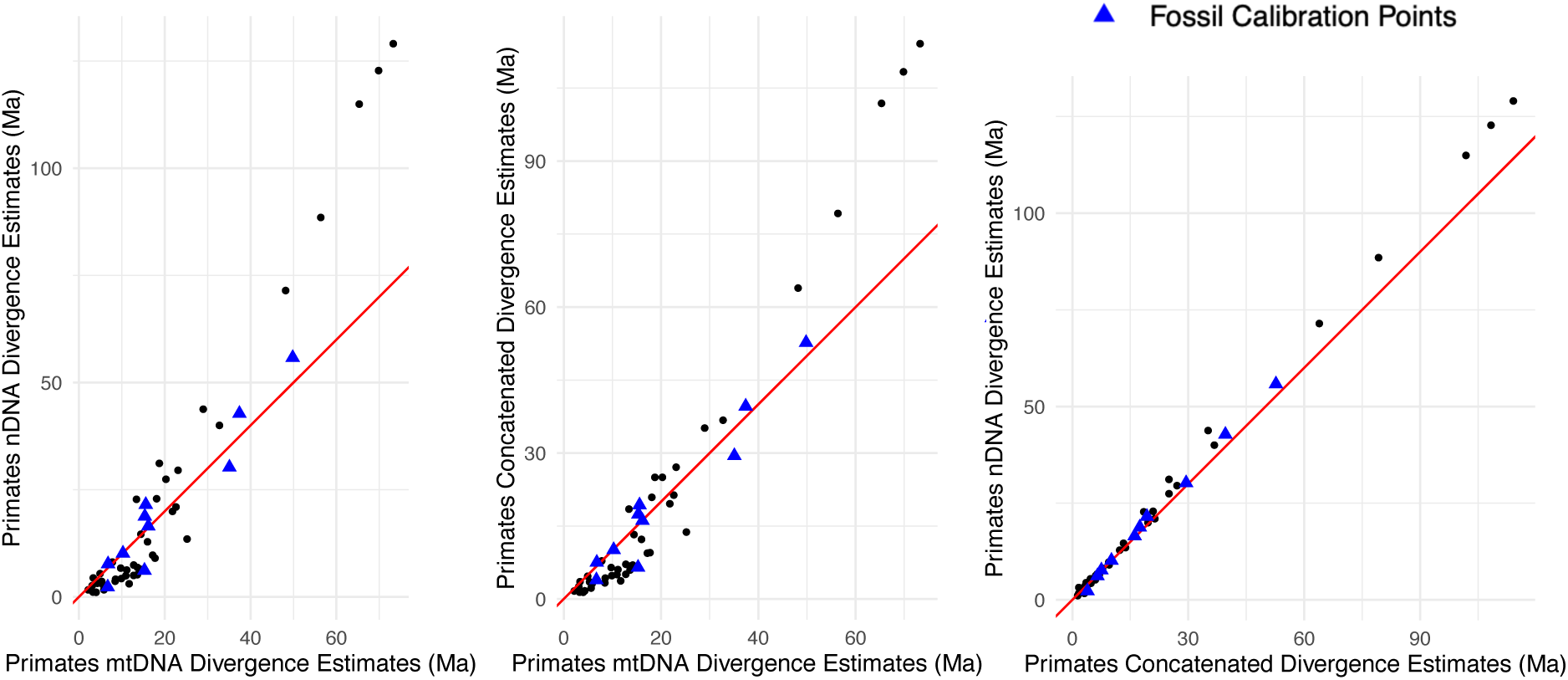
Divergence estimates for mtDNA, nDNA and concatenated runs compared in 3 independent plots. From left to right: (a) mtDNA dates (x) vs nDNA dates (y), (b) mtDNA dates (x) vs concatenated dates (y), and (c) concatenated dates (x) vs nDNA dates (y).

Observed and expected genetic distances for all pairwise combinations of the mtDNA and nDNA dataset for primates are plotted in Figure 2. As predicted, the mtDNA dataset showed extensive saturation for 3rd codon sites, while the nDNA dataset showed little to no saturation.

**Fig. 2.**
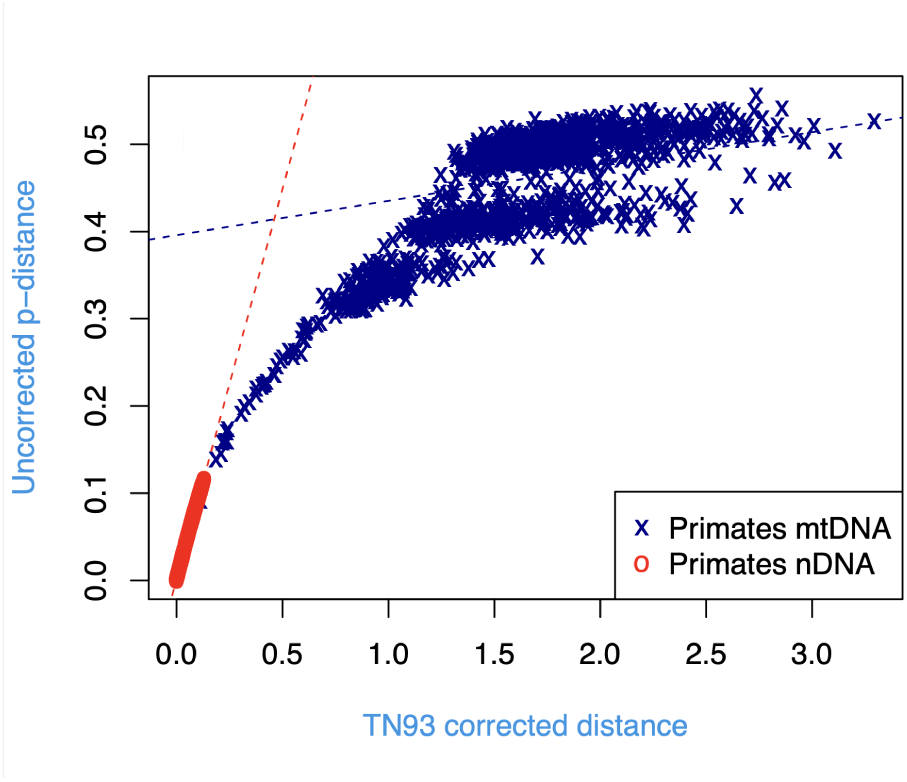
TN93 corrected distances (y) against uncorrected p-distances for all 3rd codon position in the mtDNA (blue) and nDNA (red) alignments. The corresponding dashed blue and red lines represent the linear regression slopes for each alignment.

The pattern of divergence discordance observed in the *Aves* dataset was generally congruent with that of primates (Fig.3), with younger nodes favouring older mtDNA estimates, and basal nodes generally falling above the line towards older nDNA/concatenated estimates. The concatenated estimates again skewed towards the nDNA dates, especially for basal nodes.

**Fig. 3.**
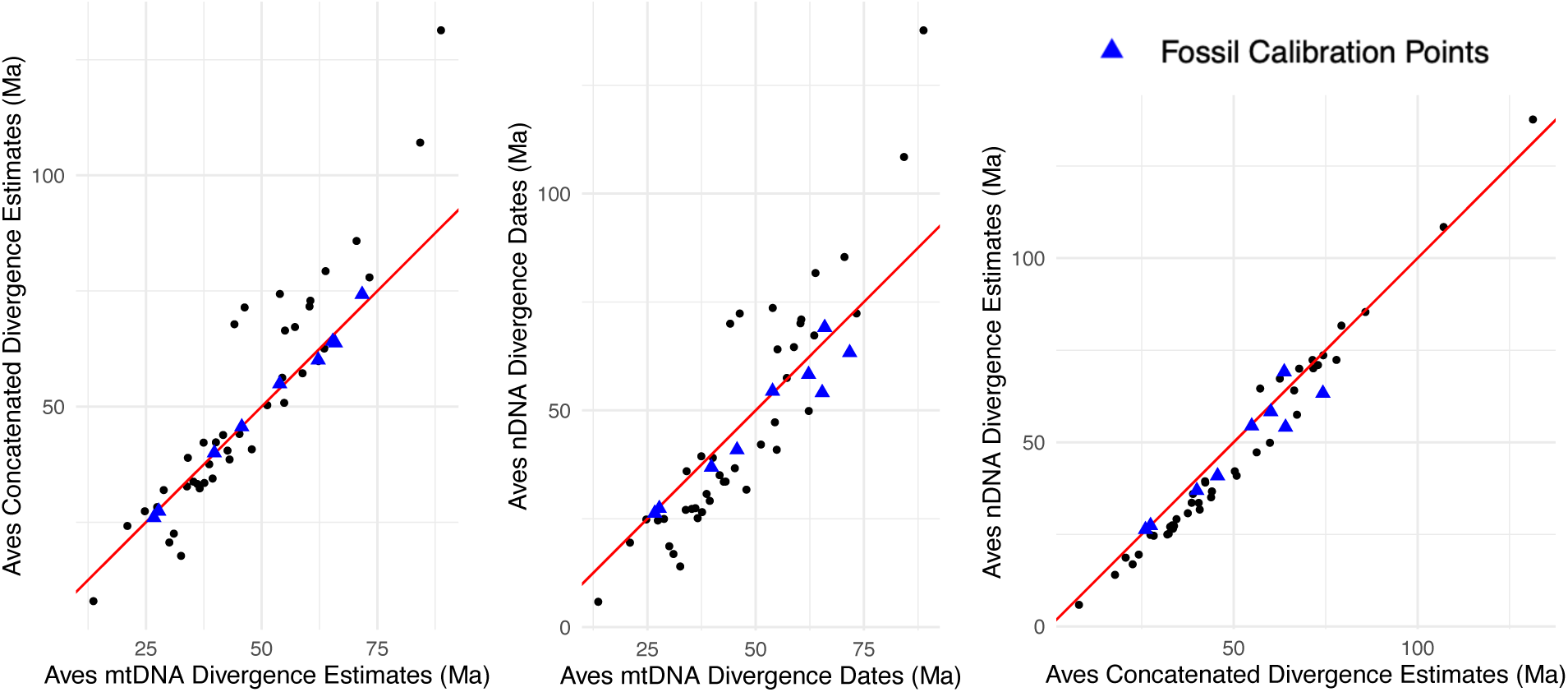
Divergence estimates for mtDNA, nDNA and concatenated runs compared in 3 independent plots. From left to right: (a) mtDNA dates (x) vs nDNA dates (y), (b) mtDNA dates (x) vs concatenated dates (y), and (c) concatenated dates (x) vs nDNA dates (y).

Unlike all other groups, the *Aves* datasets did not contain any species level relationships, and thus the observed and expected pairwise genetic distances for the mtDNA dataset started out significantly larger in comparison with other datasets (Fig.4). Regardless, the extent of saturation was again significantly higher in the mtDNA dataset than nDNA.

**Fig. 4.**
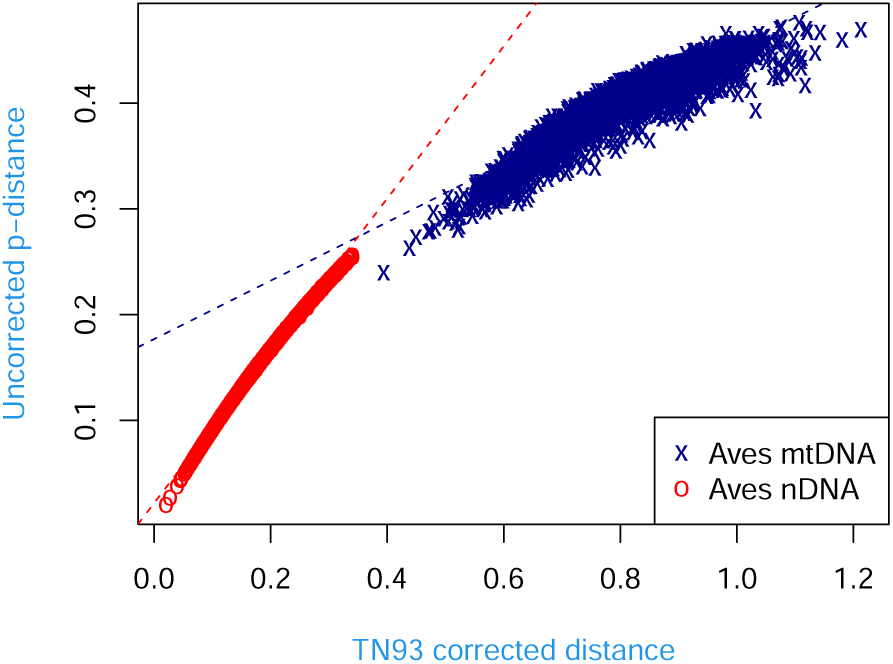
TN93 corrected distances (y) against uncorrected p-distances for all 3rd codon position in the mtDNA (blue) and nDNA (red) alignments. The correseponding dashed blue and red lines represent the linear regression slopes for each alignment.

The trend of divergence discordance for *Squamata* was congruent with *Primates* and *Aves* even though their crown ages are vastly different (Fig.5). Squamate mtDNA estimates are older for younger nodes and nDNA estimates are older towards the root node, although the degree of nuclear skew is to a lesser extent than in the two aforementioned groups. Again, the concatenated dates are highly congruent with the nDNA dates over the mtDNA estimates (Fig. 5c).

**Fig. 5.**
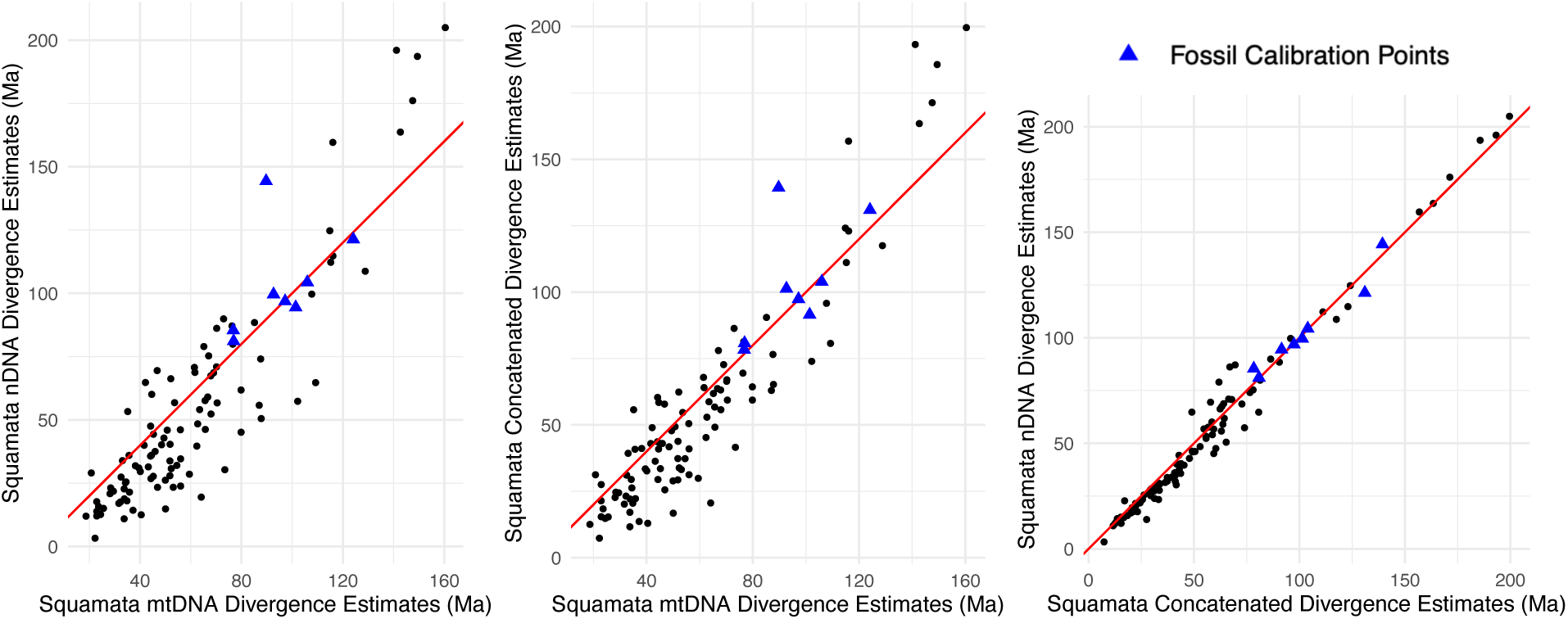
Divergence estimates for mtDNA, nDNA and concatenated runs compared in 3 independent plots. From left to right: (a) mtDNA dates (x) vs nDNA dates (y), (b) mtDNA dates (x) vs concatenated dates (y), and (c) concatenated dates (x) vs nDNA dates (y).

### Substitution saturation

The *Squamata* datasets again show congruent results with the other groups, despite larger overall genetic distances overall due to an older crown age (Fig.6). The mtDNA alignment is saturated as hypothesised, unlike the nDNA alignment.

**Fig. 6.**
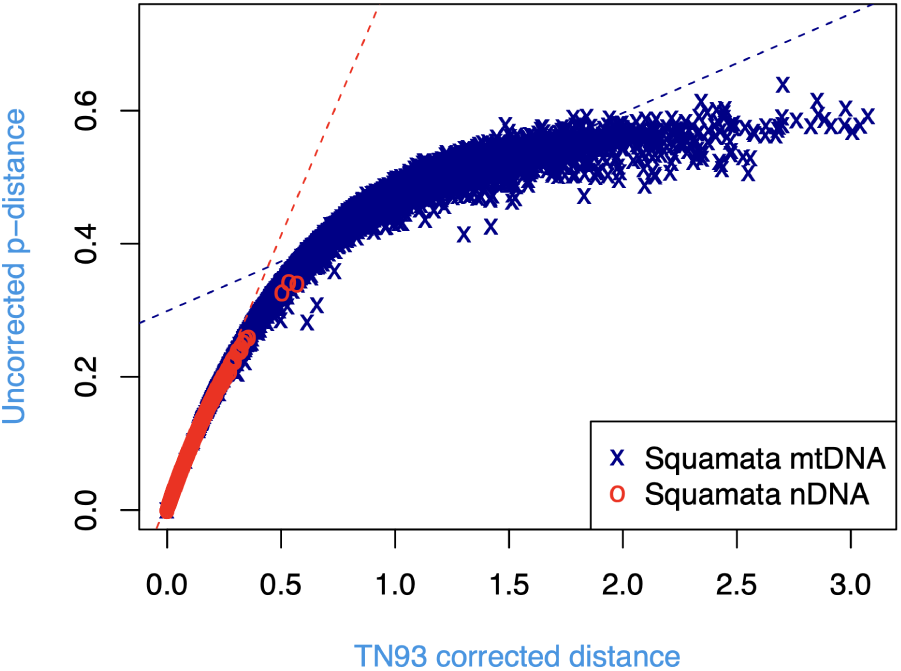
TN93 corrected distances (y) against uncorrected p-distances for all 3rd codon position in the mtDNA (blue) and nDNA (red) alignments. The correseponding dashed blue and red lines represent the linear regression slopes for each alignment.

In contrast to the other groups, the anuran datasets revealed an opposing skew in terms of basal node divergences (Fig.7), with slightly older mitochondrial estimates for crown nodes. However, shallower nodes still reported older mtDNA estimates, and concatenated matrix dates still skewed heavily towards the nDNA dates.

**Fig. 7.**
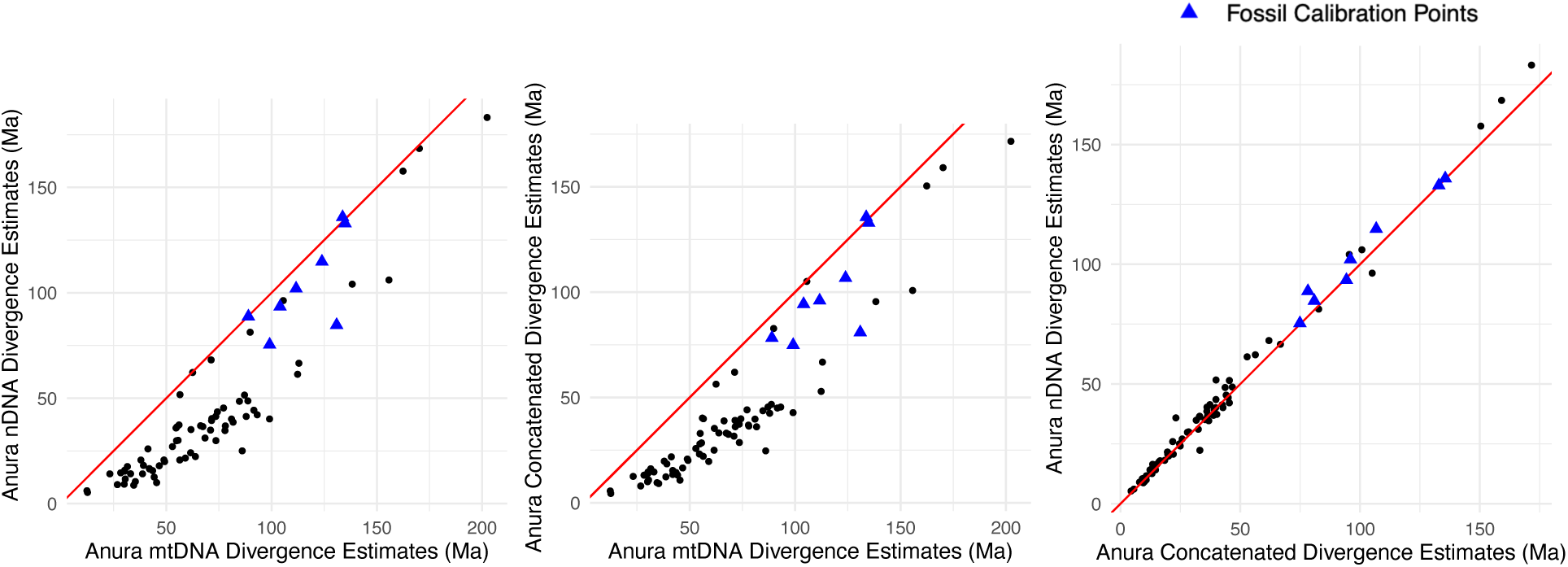
Divergence estimates for mtDNA, nDNA and concatenated runs compared in 3 independent plots. From left to right: (a) mtDNA dates (x) vs nDNA dates (y), (b) mtDNA dates (x) vs concatenated dates (y), and (c) concatenated dates (x) vs nDNA dates (y).

Interestingly, the anuran nDNA dataset displayed substitution saturation (Fig.8), as well as the least disparity between mtDNA and nDNA regression slopes, indicating a similar extent of saturation between the datasets, a surprising result considering the large disparity between nDNA and mtDNA saturation observed in all other tetrapod groups.

**Fig. 8.**
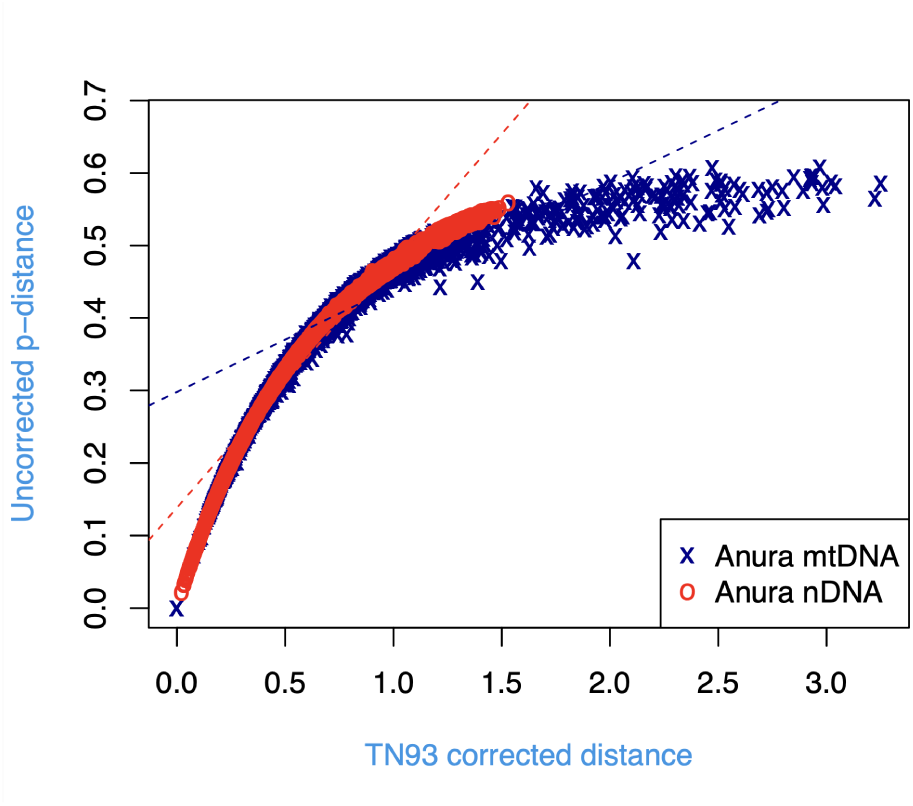
TN93 corrected distances (y) against uncorrected p-distances for all 3rd codon position in the mtDNA (blue) and nDNA (red) alignments. The corresponding dashed blue and red lines represent the linear regression slopes for each alignment.

## Discussion

### Congruent patterns of discordance across tetrapods

Nearly all groups displayed the expected trend of older mtDNA ages for younger nodes, and older nDNA estimates for basal nodes, with the only exception being for the anuran datasets, wherein basal node ages were moderately older for the mtDNA estimates.

Past studies exploring mito-nuclear divergence discordance have been limited in terms of taxon sampling and calibration strategies. For example, Arbogast et al. (2002) investigated the theoretical distribution of divergence dates from saturated data around just a single fixed calibration prior, and Zheng et al. (2011) employed a prior constraining the crown node to understand the distribution of mtDNA and nDNA estimates throughout the phylogenetic tree for salamanders. In contrast, this study investigates trends of discordance across a wider range of taxonomic groups while employing log-normal distributions to a range of calibration points that constrain recent to intermediate nodes (for which fossil data is abundant), rather than adding constraints to crown nodes.

Given that the methods used in this study are extensively utilised for divergence dating, this general trend of overestimation of mtDNA dates for shallow nodes and underestimation of basal node ages is likely to be the prevailing pattern across all tetrapod phylogenies, regardless of crown age (supposing that the mtDNA alignment is saturated and the nDNA alignment is not). This trend is exemplified by the pattern observed between primates/birds and squamates, two groups with significantly contrasting crown ages which display congruent patterns of discordance.

Interestingly, the timing of the switchover from older mtDNA ages for younger nodes to older nDNA estimates for deeper nodes is not comparable across groups. For primates, the trend of older nDNA dates begins at shallower nodes than for other groups (relative to their respective crown ages). Apart from the obvious difference in crown ages between the groups, this difference in crossover points from older mtDNA to older nDNA estimates is also likely dictated by the differing distributions of calibration points for each group. For example, for the primate datasets, the available calibration points constrained several recent genus-level relationships (since this was the range wherein fossil data was most abundant), while fossil calibrations for the other groups constrained a number of intermediate nodes. Since dates from saturated data become skewed towards fossil calibration points (Arbogast et al., 2002; Zheng et al., 2011), this difference in fossil placement is likely to have caused the more recent crossover into older nDNA estimates (∼ 15 − 20 Ma) for primates compared to the other groups. However, for the anuran datasets, the mtDNA dates remained older throughout the tree, although this result likely has more to do with substitution saturation than calibration distributions. Nonetheless, this finding highlights the importance of fossil calibration placement in influencing the trend of divergence dates estimated using saturated data.

It is also worth noting that for primates and birds, the crown ages obtained are substantially older for nDNA than any estimates from prior dating studies. However, (Perelman et al., 2011) constrained the crown node of primates to be no older than 90 Ma, and (Hackett et al., 2008) never used their alignment for divergence dating. Additionally, as this study’s intention is not to obtain accurate dates in line with other studies, but rather to understand discordance after controlling for *a priori* parameters, these rather lofty dates for nDNA crown nodes have little bearing on the general pattern obtained at the end of the MCMC analyses.

### A note on divergence dating using Order Anura

The results obtained for the anuran datasets break the pattern observed in all other tetrapod groups thus far. While recent divergences are overestimated by mtDNA as in other groups, basal node ages are marginally older for the mtDNA dataset compared to nDNA. Additionally, the anuran saturation plots were unique, as the nDNA dataset also displayed substitution saturation, unlike in the other tetrapod groups. As both datasets displayed substitution saturation to similar extents, basal node ages for both trees were similarly affected by saturation, and thus their divergence estimates appeared more or less congruent. Importantly, despite sharing similar clade ages with anurans, the squamate datasets did not display nDNA saturation whatsoever, indicating that the lack divergence discordance in the anuran dataset is particularly unique among tetrapods and not entirely influenced by crown age.

The substitution saturation in the anuran nDNA dataset could well be an artefact of the dataset used, since Feng et al. (2017) also attested that their crown age was younger than previous estimates for the group across differing data types. Nonetheless, this suggests that mtDNA and nDNA datasets that undergo saturation can provide congruent yet equally misleading estimates for basal divergence times. Furthermore, these results emphasize the need to check for saturation prior to divergence dating, a conclusion that is not unique (Farias et al., 2001; Roje, 2010; Brabec et al., 2012), but one that must also apply to large nDNA datasets that cover a wide range of taxa, as they may not always be exempt from the effects of saturation bias (Duchêne et al., 2022).

### Heavy bias in estimates using mito-nuclear concatenation

Despite controlling for the number of variable sites in each alignment, estimates from concatenation in all groups skewed heavily towards the nDNA estimates. It is interesting to note that even for anurans, a group that did not display significantly differing saturation curves between mtDNA and nDNA, divergence dates were still skewed towards the nDNA estimates, indicating that saturation alone may not explain this nDNA bias.

In our initial analyses of concatenated matrices for primates, each gene and codon position in the mtDNA alignment was allotted a separate partition, while only 3 nDNA partitions were used, one for each codon position. Furthermore, variable sites were not controlled for, and nDNA variable sites far outnumbered those of the mtDNA alignment. Yet, the dates estimated using the concatenated dataset skewed heavily towards the mtDNA dates (see Supplemental File S2), a surprising contrast to our final results. This could indicate that differential partitioning schemes may also play a role in dictating the direction of skew for the date estimates of concatenated alignments, although this aspect needs to be tested further.

Regardless of the direction of skew, it is clear that dating concatenated alignments entails a heavy estimation bias towards mtDNA or nDNA. As most current supermatrix phylogenies use comparable coverages of mtDNA and nDNA markers (Jønsson et al., 2016; McCraney et al., 2020; Henríquez-Piskulich et al., 2024), the results of such analyses would likely yield a heavy nDNA skew of divergence estimates, as displayed in all groups investigated in this study. As a result, one could make a strong case for the utilisation of mtDNA data in alignments that lack a large number of parsimony-informative sites, since mtDNA markers can increase the amount of phylogenetic information available in the alignment without introducing an inherent bias to the final date estimates. However, if one aims to assemble a dated phylogeny using a concatenated matrix, it is imperative that the mtDNA and nDNA alignments contain comparable amounts of missing data and variable sites, while employing a balanced partitioning scheme with approximately the same number of mtDNA and nDNA partitions. Failure to adhere to any of the aforementioned criteria might still introduce a mtDNA skew to the final estimates.

### Accounting for pitfalls in nDNA datasets

Other than the anuran datasets, all tetrapod groups investigated in this study displayed congruent patterns of discordance corresponding to increased levels of mtDNA saturation compared to nDNA. However, this result does not particularly lend support to the credibility of nDNA markers in generating reliable divergence estimates, but rather flags the issue of saturation as a cause for unreliable estimation irrespective of the type of marker employed. While it is true that alignments comprising a multitude of nuclear genetic markers generally increase resolution across the phylogenetic tree, these large nDNA datasets have their respective pitfalls when it comes to employing molecular clock methods, apart from just the saturation observed in the anuran nDNA dataset. One potential hindrance is known as the ‘power gap’ effect (Wilke et al., 2009), a phenomenon that causes an overall underestimation of divergence estimates due to a lack of evidence (and thus increased stochasticity) for resolving recent divergences <10 Ma, yet is rarely brought up as a reason for unreliable divergence estimation in literature. Nuclear markers are undoubtedly more prone to the effects of the power gap due to their slow substitution rate, and thus lack of informative power for resolving shallow nodes. The primate datasets used in this study in particular contained a plethora of species-level relationships (eg. *Pan paniscus* and *Pan troglodytes*), significantly more than the other tetrapod datasets. Thus, it is plausible that the dates inferred from Fig. 1a for recent divergences could well be caused by nDNA underestimation due to the power gap effect, rather than the observed mtDNA ‘overestimates’ for shallow nodes.

Ultimately, this study advocates for the need to test for saturation in alignments regardless of data type and timescale before employing molecular dating methods, in order to avoid overestimating recent divergence times and underestimating basal divergence times. Additionally, it emphasises the need for newer substitution and clock models to mitigate the effects of saturation across phylogenetic trees, rather than advocating for the use of a particular data type for future divergence dating studies (although the use of mtDNA is still cautioned due to its rapidly evolving nature). Lastly, this study highlights the inherent nDNA bias of divergence estimates acquired using concatenated mito-nuclear matrices, and advocates for the use of mtDNA in such matrices given that a comparable protocol is followed.

## Notes

### Competing Interest Statement

The authors have declared no competing interest.

