## Supplementary File S1 for "Conflicting Timelines: Exploring patterns of mito-nuclear discordance in divergence estimates among tetrapods"

**Supplementary File S1: Taxon sampling**

Order *Primates*

Taxa used for mtDNA alignment:

*Loris tardigradus*

*Nycticebus coucang*

*Perodicticus potto*

*Galago senegalensis*

*Otolemur crassicaudatus*

*Daubentonia madagascariensis*

*Propithecus verreauxii*

*Eulemur mongoz*

*Eulemur fulvus*

*Eulemur macaco*

*Lemur catta*

*Varecia variegata*

*Tarsius bancanus*

*Tarsius syrichta*

*Aotus lemurinus*

*Cebus albifrons*

*Saimiri sciureus*

*Saguinus oedipus*

*Callicebus donacophilus*

*Ateles belzebuth*

*Colobus guereza*

*Nasalis larvatus*

*Piliocolobus badius*

*Presbytis melalophos*

*Pygathrix nemaeus*

*Semnopithecus entellus*

*Trachypithecus obscurus*

*Allenopithecus nigroviridis*

*Cercocebus torquatus*

*Cercopithecus diana*

*Cercopithecus lhoesti*

*Cercopithecus mitis*

*Chlorocebus aethiops*

*Chlorocebus sabaeus*

*Erythrocebus patas*

*Lophocebus aterrimus*

*Macaca sylvanus*

*Macaca mulatta*

*Macaca fascicularis*

*Macaca thibetana*

*Mandrillus sphinx*

*Papio hamadryas*

*Theropithecus gelada*

*Gorilla gorilla*

*Homo sapiens*

*Hylobates agilis*

*Hylobates lar*

*Nomascus siki*

*Pan paniscus*

*Pan troglodytes*

*Pongo abelii*

*Pongo pygmaeus*

*Symphalangus syndactylus*

*Galeopterus variegatus*

Exclusions/inclusions in nDNA alignment:

Since Perelman et al.’s nDNA dataset contained data for a lot more species, that dataset was pruned to include only the aforementioned species from the mtDNA list.

Class *Aves*

Taxa used for mtDNA alignment:

*Alligator mississippiensis*

*Aythya americana*

*Anser anser*

*Apteryx australis*

*Acanthisitta chloris*

*Ardea cinerea*

*Aegotheles cristatus*

*Arenaria interpres*

*Alectura lathami*

*Anas platyrhynchos*

*Anhinga rufa*

*Anseranas semipalmata*

*Buteo buteo*

*Bombycilla cedrorum*

*Bucorvus leadbeateri*

*Balaeniceps rex*

*Casuarius casuarius*

*Crotophaga ani*

*Cathartes aura*

*Ciconia ciconia*

*Corvus cornix*

*Caprimulgus indicus*

*Coturnix japonica*

*Colinus virginianus*

*Columba livia*

*Cacatua moluccensis*

*Cuculus poliocephalus*

*Crax rubra*

*Centropus sinensis*

*Diomedea chrysostoma*

*Dromaius novaehollandiae*

*Dryocopus pileatus*

*Eudromia elegans*

*Eudyptula minor*

*Eudocimus ruber*

*Fringilla coelebs*

*Falco peregrinus*

*Fregata sp.*

*Geococcyx californianus*

*Grus canadensis*

*Gallus gallus*

*Gavia stellata*

*Geotrygon violacea*

*Haematopus ater*

*Heliornis fulica*

*Jacana jacana*

*Larus vegae*

*Micrastur gilvicollis*

*Malurus melanocephalus*

*Menura novaehollandiae*

*Mionectes oleagineus*

*Morus serrator*

*Nyctibius grandis*

*Numida meleagris*

*Phodilus badius*

*Phalacrocorax carbo*

*Pelecanus conspicillatus*

*Podiceps cristatus*

*Psittacus erithacus*

*Picathartes gymnocephalus*

*Pandion haliaetus*

*Phaethornis hispidus*

*Phaethon lepturus*

*Passer montanus*

*Pitta nympha*

*Phoenicopterus ruber*

*Rhea americana*

*Regulus calendula*

*Rhynochetos jubatus*

*Sylvia atricapilla*

*Struthio camelus*

*Strix leptogrammica*

*Scytalopus magellanicus*

*Sagittarius serpentarius*

*Smithornis sharpei*

*Tyto alba*

*Tinamus guttatus*

*Turdus hortulorum*

*Thamnophilus nigrocinereus*

*Trogon viridis*

*Upupa epops*

*Vidua chalybeata*

Exclusions/inclusions in nDNA alignment:

The mtDNA alignment here was pruned to contain taxa that belonged to the same genera (preferably species) as the nDNA alignment. Regardless, there were some species differences (of taxa belonging to the same genera) between the mtDNA and nDNA alignments, as follow:

• *Aegotheles insignis*

*• Anser erythropus*

*• Ardea herodias*

*• Bombycilla garrulus*

*• Bucorvus abyssinicus*

*• Buteo jamaicensis*

*• Caprimulgus longirostris*

*• Centropus viridis*

*• Colinus cristatus*

*• Corvus corone*

*• Coturnix coturnix*

*• Crax alector*

*• Crotophaga sulcirostris*

*• Cuculus canorus*

*• Diomedea nigripes*

*• Eudocimus albus*

*• Falco mexicanus*

*• Fregata magnificens*

*• Fringilla montifringilla*

*• Gavia immer*

*• Geotrygon montana*

*• Haematopus ostralegus*

*• Larus marinus*

*• Micrastur semitorquatus*

*• Mionectes macconnelli*

*• Morus bassanus*

*• Pelecanus occidentalis*

*• Phaethornis griseogularis*

*• Podiceps auritus*

*• Pitta guajana*

*• Sylvia nana*

*• Turdus falklandii*

*• Trogon personatus*

Order *Squamata*

Taxa used for mtDNA alignment:

*Acanthosaura lepidogaster*

*Acontias meleagris*

*Acrochordus granulatus*

*Agama agama*

*Agkistrodon contortrix*

*Amphiesma stolatum*

*Amphiglossus splendidus*

*Amphisbaena fuliginosa*

*Anelytropsis papillosus*

*Anilius scytale*

*Anniella pulchra*

*Anolis carolinensis*

*Aparallactus werneri*

*Aspidites melanocephalus*

*Aspidoscelis tigris*

*Azemiops feae*

*Bipes biporus*

*Bipes canaliculatus*

*Boa constrictor*

*Bothrops asper*

*Brachylophus fasciatus*

*Brachymeles gracilis*

*Brookesia brygooi*

*Calabaria reinhardtii*

*Calotes emma*

*Casarea dussumieri*

*Causus defilippii*

*Chalarodon madagascariensis*

*Chamaeleo calyptratus*

*Chlamydosaurus kingii*

*Coleonyx variegatus*

*Colobosaura modesta*

*Coluber constrictor*

*Cordylosaurus subtessellatus*

*Cordylus namaquensis*

*Cricosaura typica*

*Crotaphytus collaris*

*Cylindrophis ruffus*

*Daboia russelii*

*Delma borea*

*Diadophis punctatus*

*Dibamus novaeguineae*

*Diplometopon zarudnyi*

*Dipsosaurus dorsalis*

*Draco blanfordii*

*Elgaria multicarinata*

*Enyalioides laticeps*

*Eublepharis macularius*

*Eugongylus rufescens*

*Eumeces schneideri*

*Exiliboa placata*

*Feylinia polylepis*

*Gambelia wislizenii*

*Gekko gecko*

*Geocalamus acutus*

*Gonatodes albogularis*

*Heloderma horridum*

*Heloderma suspectum*

*Homalopsis buccata*

*Hydrosaurus amboinensis*

*Imantodes cenchoa*

*Lacerta viridis*

*Lachesis muta*

*Lampropeltis getula*

*Lamprophis fuliginosus*

*Lanthanotus borneensis*

*Laticauda colubrina*

*Leiocephalus barahonensis*

*Leiolepis belliana*

*Leiosaurus catamarcensis*

*Lepidophyma flavimaculatum*

*Leptotyphlops humilis*

*Lialis burtonis*

*Lichanura trivirgata*

*Liolaemus bellii*

*Liolaemus elongatus*

*Liotyphlops albirostris*

*Loxocemus bicolor*

*Lycophidion capense*

*Micrurus fulvius*

*Naja kaouthia*

*Natrix natrix*

*Notechis scutatus*

*Ophisaurus attenuatus*

*Ophisaurus ventralis*

*Oplurus cyclurus*

*Pareas hamptoni*

*Petrosaurus mearnsi*

*Phelsuma lineata*

*Pholidobolus macbrydei*

*Phrynocephalus mystaceus*

*Phrynosoma platyrhinos*

*Phymaturus palluma*

*Physignathus cocincinus*

*Physignathus lesueurii*

*Plestiodon fasciatus*

*Plestiodon skiltonianus*

*Pogona vitticeps*

*Polychrus marmoratus*

*Pristidactylus torquatus*

*Python molurus*

*Rhacodactylus auriculatus*

*Rhineura floridana*

*Saltuarius cornutus*

*Sauromalus ater*

*Sceloporus variabilis*

*Scincus scincus*

*Shinisaurus crocodilurus*

*Sonora semiannulata*

*Sphenodon punctatus*

*Sphenomorphus solomonis*

*Stenocercus guentheri*

*Strophurus ciliaris*

*Takydromus sexlineatus*

*Teius teyou*

*Teratoscincus scincus*

*Thamnophis marcianus*

*Tiliqua scincoides*

*Trachyboa boulengeri*

*Trachylepis quinquetaeniata*

*Trapelus agilis*

*Trimorphodon biscutatus*

*Trogonophis wiegmanni*

*Tropidophis haetianus*

*Tupinambis teguixin*

*Uma scoparia*

*Ungaliophis continentalis*

*Uranoscodon superciliosus*

*Uromastyx aegyptia*

*Uropeltis melanogaster*

*Uta stansburiana*

*Varanus acanthurus*

*Varanus exanthematicus*

*Varanus salvator*

*Xantusia vigilis*

*Xenochrophis piscator*

*Xenodermus javanicus*

*Xenopeltis unicolor*

*Zonosaurus ornatus*

Exclusions/inclusions in nDNA alignment:

The nDNA alignment was pruned to include almost all the same taxa to species level, although the following five were also included since they fall within the same genera as taxa from the mtDNA alignment:

• *Cordylus sp*

*• Hydrosaurus sp*

*• Ophisaurus sp*

*• Sauromalus sp*

*• Takydromus sp*

Order *Anura*

Taxa used for mtDNA alignment:

*Agalychnis callidryas*

*Agalychnis lemur*

*Aglyptodactylus madagascariensis*

*Alsodes gargola*

*Alytes obstetricans*

*Amazophrynella minuta*

*Amolops loloensis*

*Amolops ricketti*

*Anaxyrus canorus*

*Anaxyrus punctatus*

*Andrias davidianus*

*Anodonthyla boulengerii*

*Aplastodiscus perviridis*

*Arthroleptis poecilonotus*

*Ascaphus truei*

*Astylosternus diadematus*

*Batrachuperus sp.*

*Batrachyla leptopus*

*Batrachyla taeniata*

*Bombina fortinuptialis*

*Bombina orientalis*

*Boophis madagascariensis*

*Brachytarsophrys feae*

*Buergeria oxycephala*

*Bufo gargarizans*

*Callulina kreffti*

*Callulops wilhelmanus*

*Calyptocephalella gayi*

*Ceratophrys cornuta*

*Chiasmocleis ventrimaculata*

*Craugastor augusti*

*Craugastor fitzingeri*

*Crinia signifera*

*Cryptobatrachus boulengeri*

*Discoglossus pictus*

*Duttaphrynus melanostictus*

*Dyscophus antongilii*

*Elachistocleis ovalis*

*Eleutherodactylus planirostris*

*Eupsophus calcaratus*

*Fejervarya limnocharis*

*Gallus gallus*

*Gastrophryne olivacea*

*Gastrotheca pseustes*

*Gastrotheca weinlandii*

*Heleophryne purcelli*

*Hemisus marmoratus*

*Hoplobatrachus tigerinus*

*Hyla chinensis*

*Hyloxalus jacobuspetersi*

*Hymenochirus boettgeri*

*Ichthyophis bannanicus*

*Incilius nebulifer*

*Insuetophrynus acarpicus*

*Kalophrynus pleurostigma*

*Kaloula conjuncta*

*Kaloula pulchra*

*Kurixalus odontotarsus*

*Leiopelma hochstetteri*

*Lepidobatrachus sp.*

*Leptobrachium chapaense*

*Leptodactylus albilabris*

*Limnodynastes salmini*

*Limnonectes fujianensis*

*Lithodytes lineatus*

*Mantophryne lateralis*

*Melanophryniscus stelzneri*

*Microhyla heymonsi*

*Mixophyes coggeri*

*Nyctimystes kubori*

*Odontophrynus occidentalis*

*Odorrana schmackeri*

*Ophryophryne microstoma*

*Oreolalax jingdongensis*

*Osteocephalus taurinus*

*Otophryne pyburni*

*Paradoxophyla palmata*

*Pelobates syriacus*

*Pelodytes ibericus*

*Pelophylax nigromaculatus*

*Peltophryne peltocephala*

*Petropedetes euskircheni*

*Phrynomantis microps*

*Phyllomedusa tomopterna*

*Physalaemus cuvieri*

*Pipa parva*

*Pipa pipa*

*Platypelis tuberifera*

*Pleurodema somuncurense*

*Pleurodema thaul*

*Polypedates megacephalus*

*Pristimantis thymelensis*

*Proceratophrys boiei*

*Pseudis paradoxa*

*Quasipaa spinosa*

*Rana amurensis*

*Rana berlandieri*

*Rana draytonii*

*Rana virgatipes*

*Ranitomeya imitator*

*Rentapia hosii*

*Rhinella marina*

*Rhinoderma darwinii*

*Rhinophrynus dorsalis*

*Scaphiophryne boribory*

*Scaphiophryne marmorata*

*Scaphiopus couchii*

*Schismaderma carens*

*Scinax ruber*

*Sooglossus thomasseti*

*Spea multiplicata*

*Stereocyclops incrassatus*

*Strongylopus grayii*

*Stumpffia pygmaea*

*Sylvirana guentheri*

*Telmatobius vellardi*

*Trichobatrachus robustus*

*Xenopus epitropicalis*

*Xenopus kobeli*

Exclusions/inclusions in nDNA alignment:

The nDNA alignment was again pruned to include common species, although two that were included were from the same genera but not the same species, namely:

*• Batrachuperus sp.*

*• Lepidobatrachus sp.*

**Supplementary File S1: Fossil calibration choices**

Order *Primates*

All fossil calibrations were sourced from the original Pozzi et al. 2014 paper that was used for the mtDNA alignment. In addition to simply utilizing more calibration points than Perelman et al. 2012 (the study from which the nDNA alignment was sourced), each calibration point was given a strong justification as well as upper and lower bounds, which further facilitated our choice of calibration points from the 2014 paper. A total of 10 fossil calibration points were employed.

Class *Aves*

The choice of calibration points was quite straightforward for Class *Aves*, since the study used for the nDNA alignment did not utilise fossil calibrations, and thus the Arcones et al. 2021 paper’s calibration points were utilised. Naturally, out of the 25 available calibration points from that paper, many could not be utilised since our mtDNA alignments were pruned to have a subset of taxa to match the nDNA alignment. A total of 9 calibration points were employed.

Order *Squamata*

Fossil calibrations for squamates were sourced from the Zheng & Wiens 2015 study (which was the source for the mtDNA alignment), as again the nDNA dataset from Wiens et al. 2012 was not used to assemble a dated phylogeny. Again, certain calibration points that did not fit our final taxa list were excluded. A total of 8 calibration points were employed.

Order *Anura*

For the anuran datasets, Feng et al. 2017’s calibrations were used (source for the nDNA alignment), since each calibration point was justified using a secondary citation and each point had marked distribution bounds. Again, only calibration points that fit our final list of taxa after pruning both mtDNA and nDNA alignments were utilised. A total of 8 calibration points were employed.
