## Supplementary File S2 for "Conflicting Timelines: Exploring patterns of mito-nuclear discordance in divergence estimates among tetrapods"

**Supplementary File S2: Saturation plots for 1^st^ and 2^nd^ codon positions**

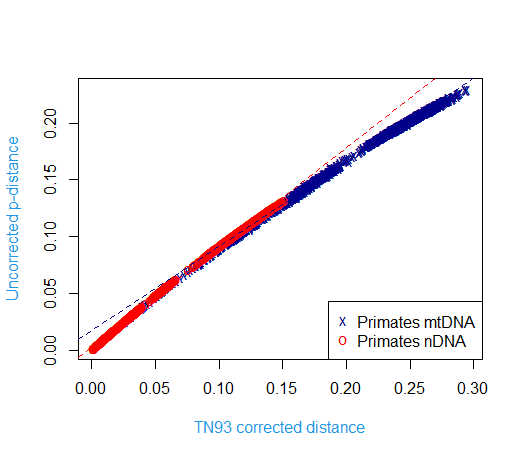
**Order *Primates*: 1^st^ codon observed vs expected genetic distances**

**Order *Primates*: 2^nd^ codon observed vs expected genetic distances**
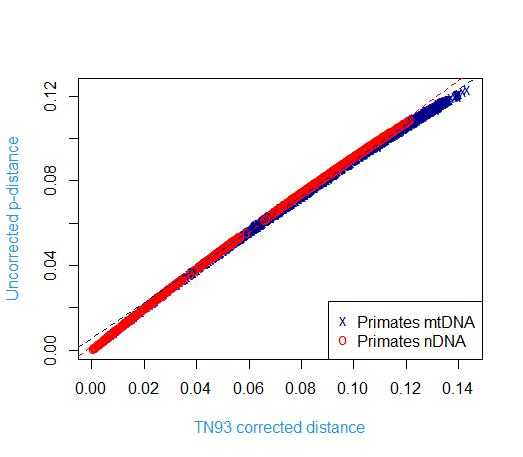

**Class *Aves*: 1^st^ codon expected vs observed genetic distances**

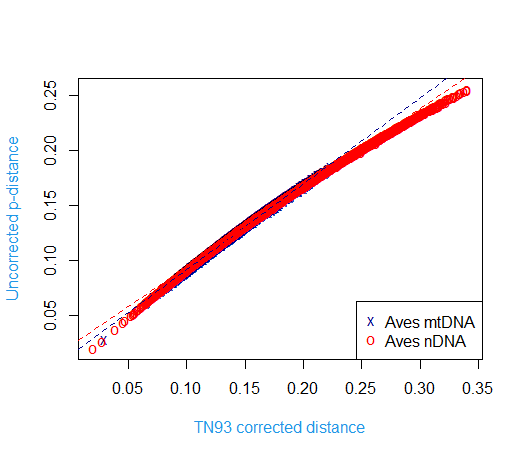

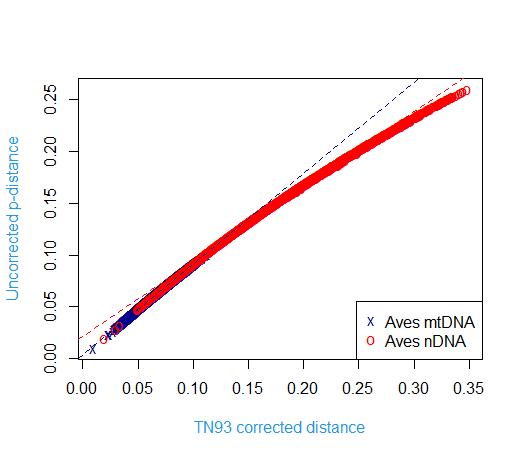
**Class *Aves*: 2^nd^ codon expected vs observed genetic distances**

**Order *Squamata*: 1^st^ codon observed vs expected genetic distances**

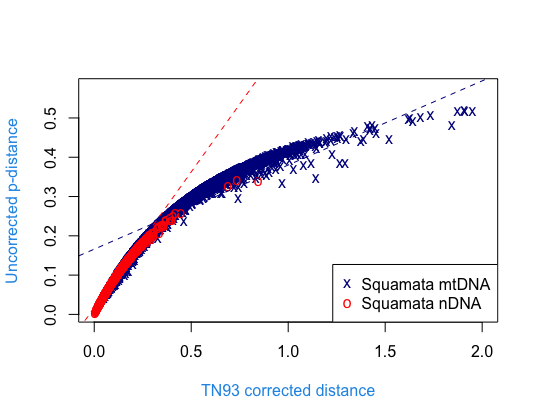

**Order *Squamata*: 2^nd^ codon observed vs expected genetic distances**

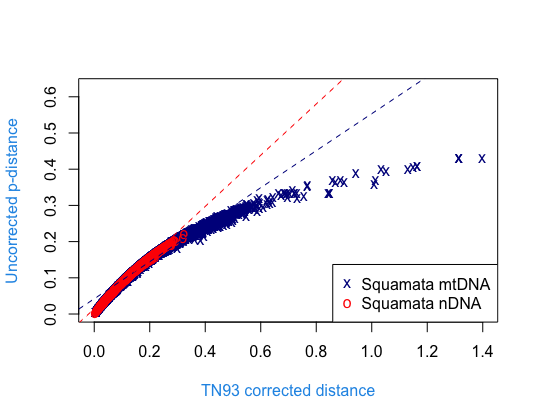

**Order *Anura*: 1^st^ codon observed vs expected genetic distances**

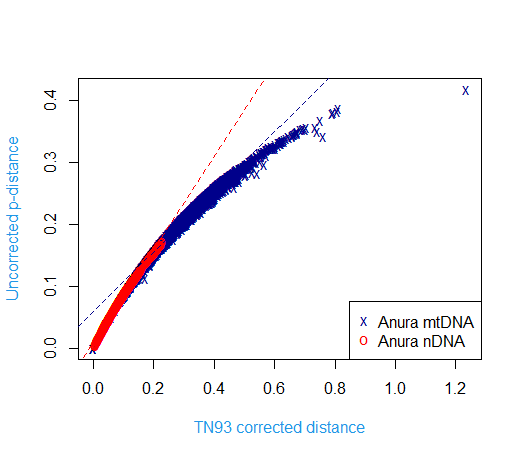

**Order *Anura*: 2^nd^ codon observed vs expected genetic distances**

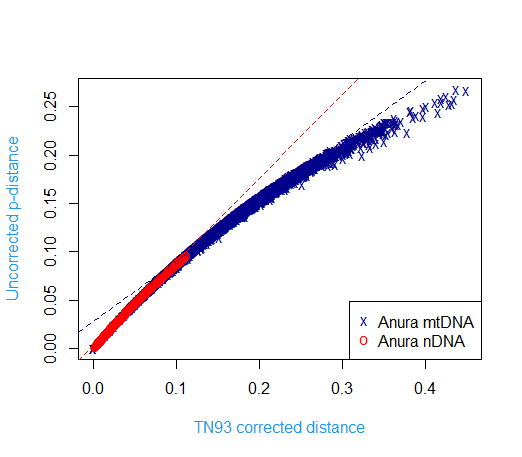

**Supplementary File S2: Divergence date estimates for all major nodes for the initial run of the mito-nuclear matrix dataset for Order *Primates***

*** Fossil calibration points**

**Mitochondrial skew of divergence estimates of the mito-nuclear matrix dataset**

| **Major Nodes** | **Nuclear (Ma) - mean crown ages** | **Mitochondrial (Ma) - mean crown ages** | **MtDNA + Nuc (Ma) - mean crown ages** |
| --- | --- | --- | --- |
| **Crown primates** | **99.48** | **70.36** | **73.58** |
| **Crown Haplorhini** | **92.02** | **62.59** | **69.20** |
| **Crown Strepsirrhini** | **75.16** | **57.08** | **61.71** |
| **Crown Lorisiformes*** | **38.14** | **38.37** | **37.50** |
| **Lemuriformes-Chiromyformes** | **62.35** | **46.82** | **52.47** |
| **Crown Lemuriformes** | **36.92** | **29.43** | **30.16** |
| **Crown Anthropoidea*** | **43.93** | **46.93** | **48.99** |
| **Crown Platyrrhini** | **24.14** | **21.65** | **23.22** |
| **Crown Catarrhini*** | **27.99** | **32.61** | **33.81** |
| **Crown Cercopithicoidea** | **13.79** | **22.28** | **23.24** |
| **Crown Colobinae** | **9.56** | **13.86** | **15.41** |
| **Crown Cercopithecinae** | **9.34** | **16.04** | **16.00** |
| **Crown Hominoidae*** | **19.28** | **21.56** | **21.75** |
| **Alleno-Cerco-Chloro clade** | **6.69** | **12.00** | **12.42** |
| **Macaque-Baboon-Mandrill clade** | **6.56** | **13.48** | **13.32** |
| **Family Homonidae*** | **16.12** | **16.17** | **16.36** |
