## Supplementary File S3 for "Conflicting Timelines: Exploring patterns of mito-nuclear discordance in divergence estimates among tetrapods"

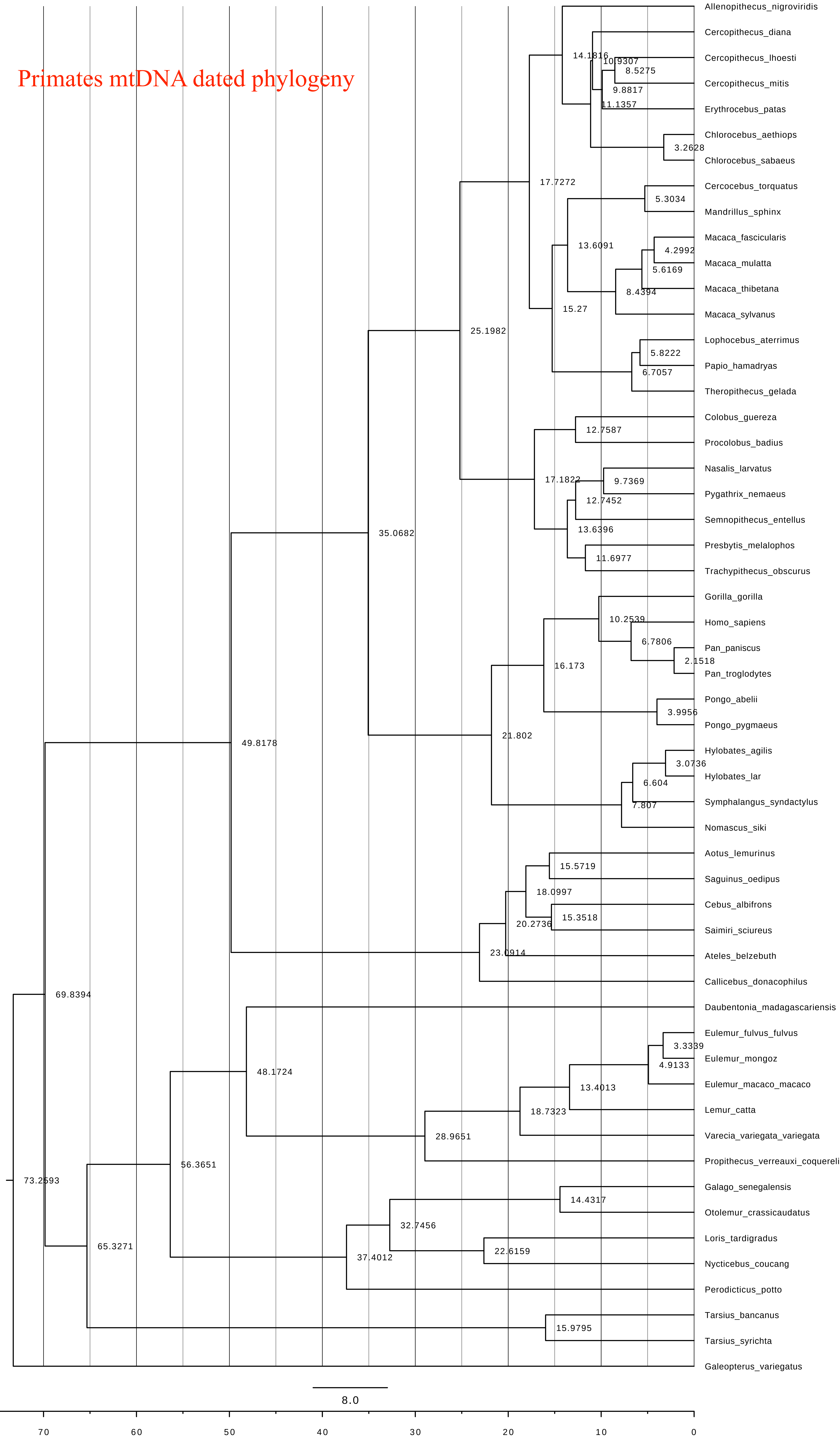

Primates mtDNA dated phylogeny
Allenopithecus_nigroviridis
Cercopithecus_diana
Cercopithecus_lhoesti
Cercopithecus_mitis
Erythrocebus_patas
Chlorocebus_aethiops
Chlorocebus_sabaeus
Cercocebus_torquatus
Mandrillus_sphinx
Macaca_fascicularis
Macaca_mulatta
Macaca_thibetana
Macaca_sylvanus
Lophocebus_aterrimus
Papio_hamadryas
Theropithecus_gelada
Colobus_guereza
Procolobus_badius
Nasalis_larvatus
Pygathrix_nemaeus
Semnopithecus_entellus
Presbytis_melalophos
Trachypithecus_obscurus
Gorilla_gorilla
Homo_sapiens
Pan_paniscus
Pan_troglodytes
Pongo_abelii
Pongo_pygmaeus
Hylobates_agilis
Hylobates_lar
Symphalangus_syndactylus
Nomascus_siki
Aotus_lemurinus
Saguinus_oedipus
Cebus_albifrons
Saimiri_sciureus
Ateles_belzebuth
Callicebus_donacophilus
Daubentonia_madagascariensis
Eulemur_fulvus_fulvus
Eulemur_mongoz
Eulemur_macaco_macaco
Lemur_catta
Varecia_variegata_variegata
Propithecus_verreauxi_coquereli
Galago_senegalensis
Otolemur_crassicaudatus
Loris_tardigradus
Nycticebus_coucang
Perodicticus_potto
Tarsius_bancanus
Tarsius_syrichta
Galeopterus_variegatus
14.1816
10.9307
8.5275
9.8817
11.1357
3.2628
17.7272
5.3034
13.6091
4.2992
5.6169
8.4394
15.27
25.1982
5.8222
6.7057
12.7587
17.1822
9.7369
12.7452
13.6396
11.6977
35.0682
10.2539
6.7806
2.1518
16.173
3.9956
21.802
49.8178
3.0736
6.604
7.807
15.5719
18.0997
15.3518
20.2736
23.0914
69.8394
48.1724
3.3339
4.9133
13.4013
18.7323
28.9651
56.3651
73.2593
14.4317
32.7456
22.6159
37.4012
65.3271
15.9795
8.0
70
60
50
40
30
20
10
0

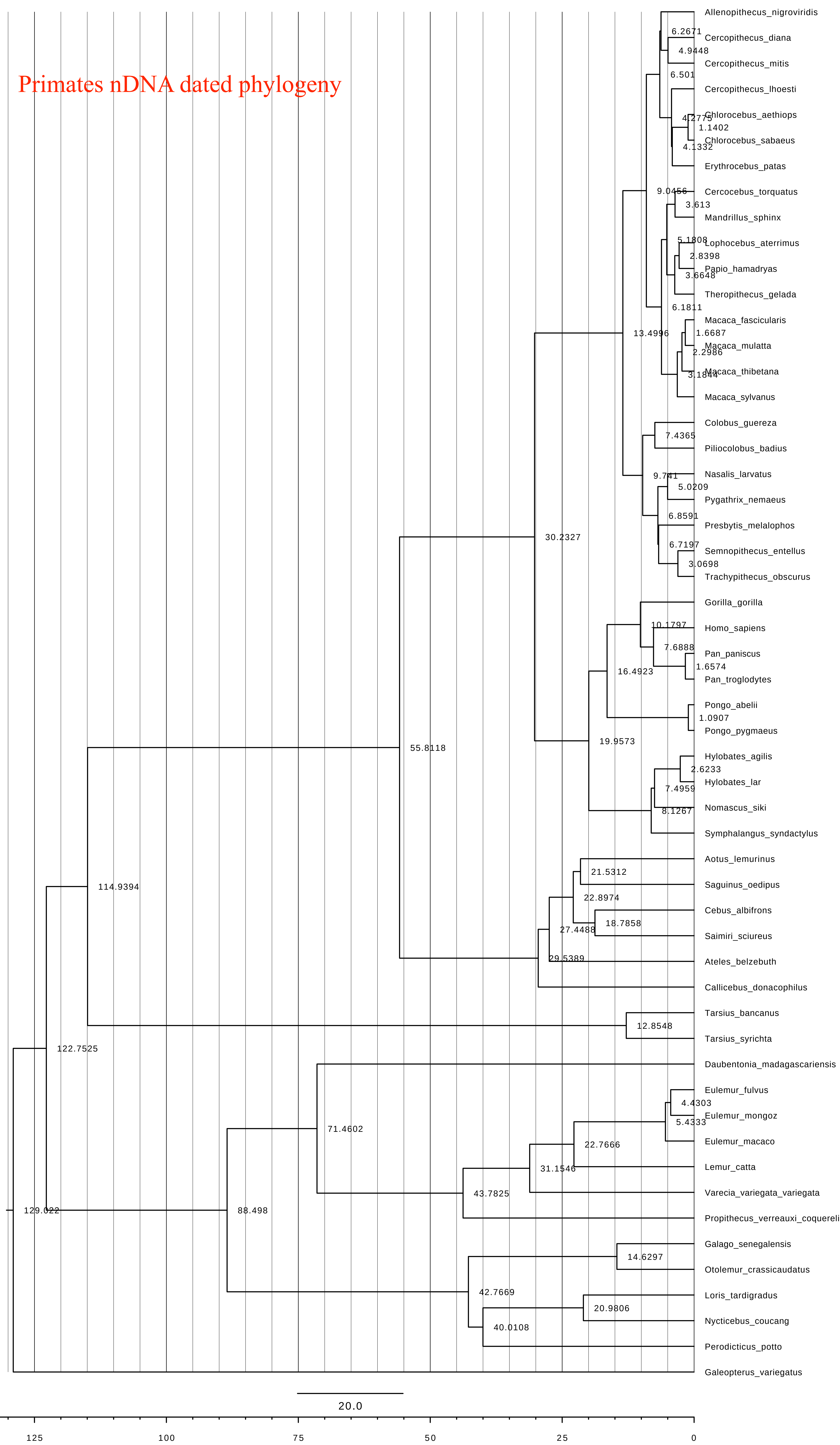
Primates nDNA dated phylogeny
Allenopithecus_nigroviridis
6.2671
Cercopithecus_diana
4.9448
Cercopithecus_mitis
6.501
Cercopithecus_lhoesti
4.2775
Chlorocebus_aethiops
1.1402
Chlorocebus_sabaeus
4.1332
Erythrocebus_patas
9.0456
Cercocebus_torquatus
3.613
Mandrillus_sphinx
5.1808
Lophocebus_aterrimus
2.8398
Papio_hamadryas
3.6648
Theropithecus_gelada
6.1811
Macaca_fascicularis
13.4996
1.6687
Macaca_mulatta
2.2986
Macaca_thibetana
3.1844
Macaca_sylvanus
Colobus_guereza
7.4365
Piliocolobus_badius
Nasalis_larvatus
9.741
5.0209
Pygathrix_nemaeus
6.8591
Presbytis_melalophos
30.2327
6.7197
Semnopithecus_entellus
3.0698
Trachypithecus_obscurus
Gorilla_gorilla
10.1797
Homo_sapiens
7.6888
Pan_paniscus
16.4923
1.6574
Pan_troglodytes
Pongo_abelii
1.0907
Pongo_pygmaeus
19.9573
55.8118
Hylobates_agilis
2.6233
Hylobates_lar
7.4959
8.1267
Nomascus_siki
Symphalangus_syndactylus
Aotus_lemurinus
21.5312
Saguinus_oedipus
114.9394
22.8974
Cebus_albifrons
18.7858
27.4488
Saimiri_sciureus
29.5389
Ateles_belzebuth
Callicebus_donacophilus
Tarsius_bancanus
12.8548
Tarsius_syrichta
122.7525
Daubentonia_madagascariensis
Eulemur_fulvus
4.4303
Eulemur_mongoz
5.4333
71.4602
Eulemur_macaco
22.7666
Lemur_catta
31.1546
43.7825
Varecia_variegata_variegata
129.022
88.498
Propithecus_verreauxi_coquereli
Galago_senegalensis
14.6297
Otolemur_crassicaudatus
42.7669
Loris_tardigradus
20.9806
Nycticebus_coucang
40.0108
Perodicticus_potto
Galeopterus_variegatus
20.0
125
100
75
50
25
0

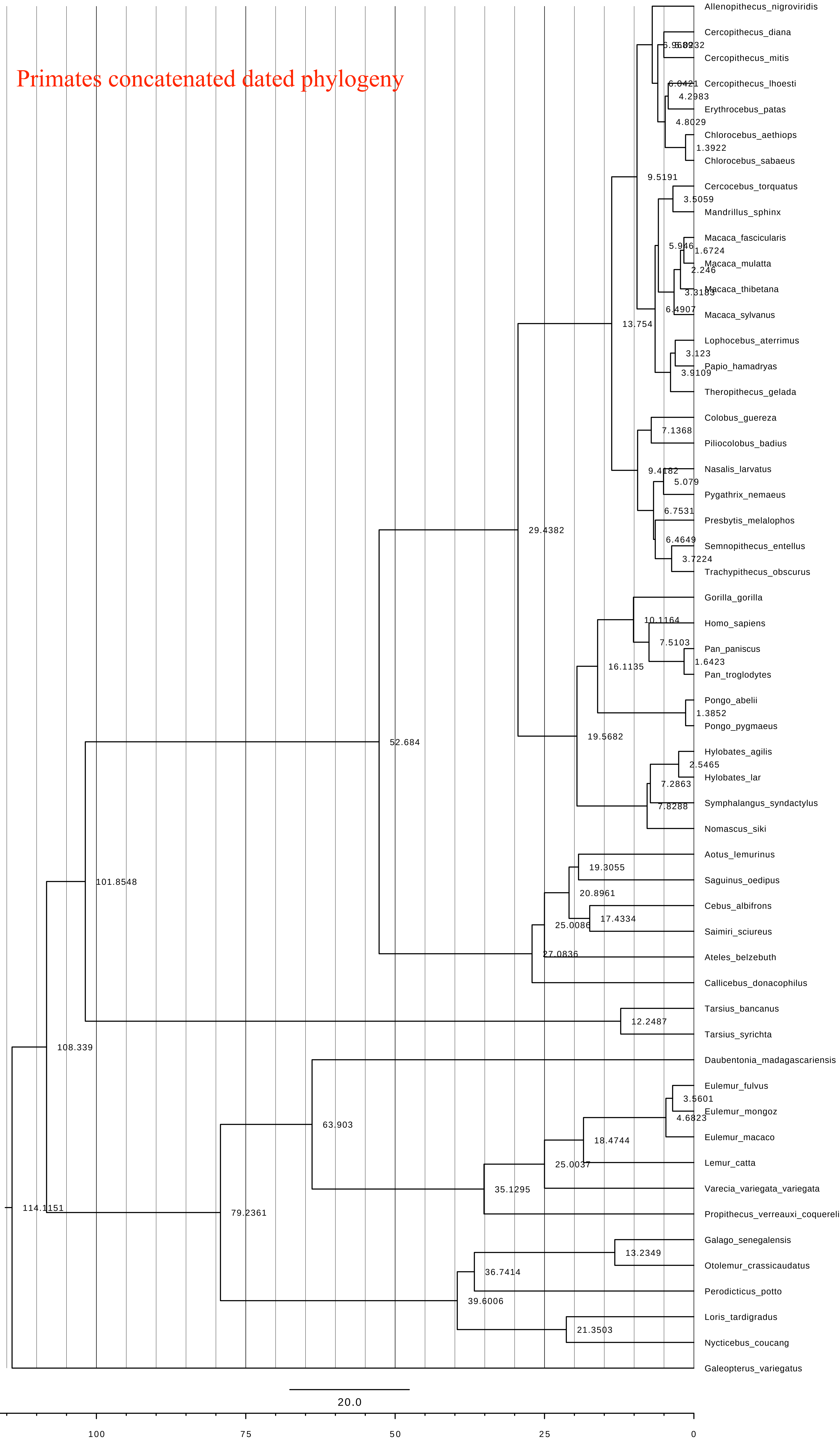
Primates concatenated dated phylogeny
Allenopithecus_nigroviridis
Cercopithecus_diana
Cercopithecus_mitis
Cercopithecus_lhoesti
Erythrocebus_patas
Chlorocebus_aethiops
Chlorocebus_sabaeus
Cercocebus_torquatus
Mandrillus_sphinx
Macaca_fascicularis
Macaca_mulatta
Macaca_thibetana
Macaca_sylvanus
Lophocebus_aterrimus
Papio_hamadryas
Theropithecus_gelada
Colobus_guereza
Piliocolobus_badius
Nasalis_larvatus
Pygathrix_nemaeus
Presbytis_melalophos
Semnopithecus_entellus
Trachypithecus_obscurus
Gorilla_gorilla
Homo_sapiens
Pan_paniscus
Pan_troglodytes
Pongo_abelii
Pongo_pygmaeus
Hylobates_agilis
Hylobates_lar
Symphalangus_syndactylus
Nomascus_siki
Aotus_lemurinus
Saguinus_oedipus
Cebus_albifrons
Saimiri_sciureus
Ateles_belzebuth
Callicebus_donacophilus
Tarsius_bancanus
Tarsius_syrichta
Daubentonia_madagascariensis
Eulemur_fulvus
Eulemur_mongoz
Eulemur_macaco
Lemur_catta
Varecia_variegata_variegata
Propithecus_verreauxi_coquereli
Galago_senegalensis
Otolemur_crassicaudatus
Perodicticus_potto
Loris_tardigradus
Nycticebus_coucang
Galeopterus_variegatus
6.0421
4.2983
4.8029
1.3922
9.5191
3.5059
5.946
1.6724
2.246
3.3183
6.4907
13.754
3.123
3.9109
7.1368
9.4182
5.079
6.7531
6.4649
3.7224
29.4382
10.1164
7.5103
1.6423
16.1135
1.3852
19.5682
2.5465
7.2863
7.8288
52.684
19.3055
20.8961
17.4334
25.0086
27.0836
101.8548
12.2487
108.339
63.903
3.5601
4.6823
18.4744
25.0037
35.1295
114.1151
79.2361
13.2349
36.7414
39.6006
21.3503
20.0
100
75
50
25
0

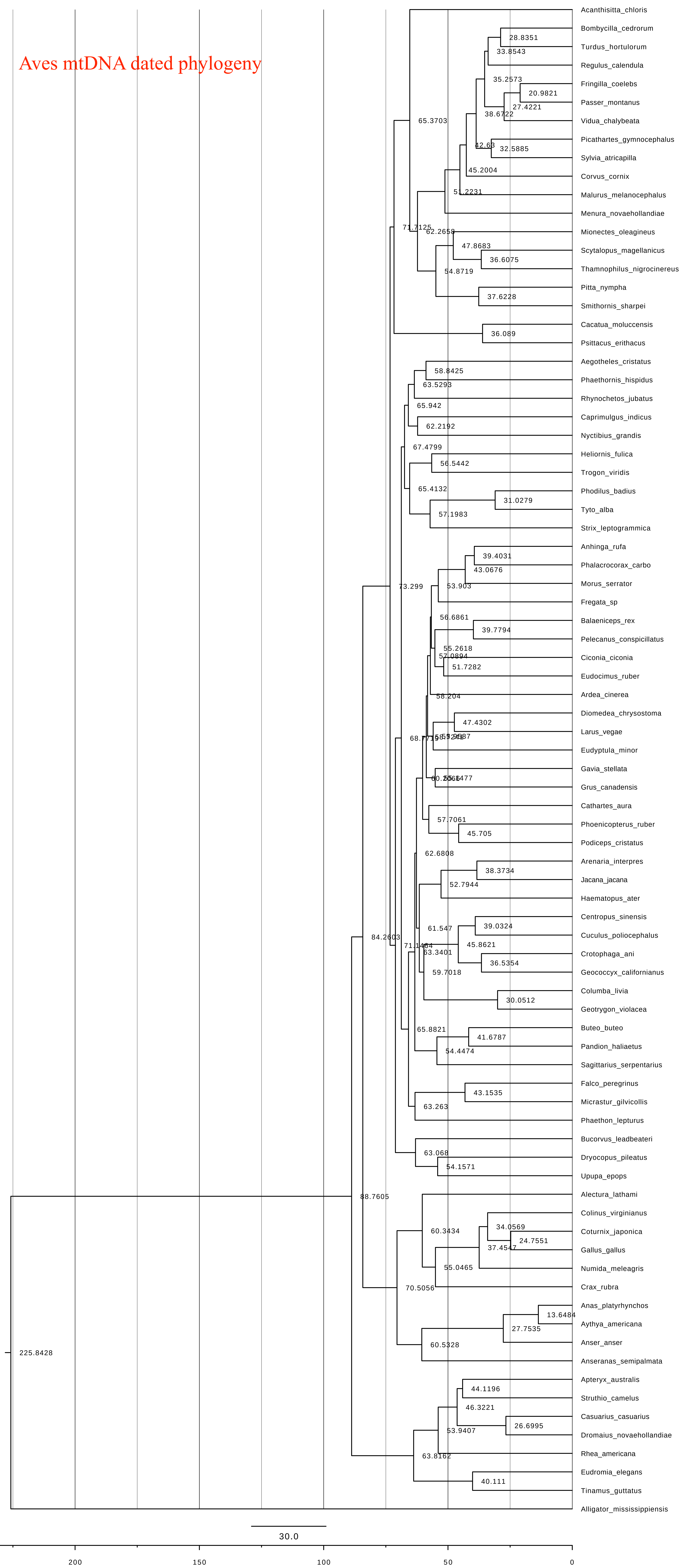

Aves mtDNA dated phylogeny
Acanthisitta_chloris
Bombycilla_cedrorum
Turdus_hortulorum
Regulus_calendula
Fringilla_coelebs
Passer_montanus
Vidua_chalybeata
Picathartes_gymnocephalus
Sylvia_atricapilla
Corvus_cornix
Malurus_melanocephalus
Menura_novaehollandiae
Mionectes_oleagineus
Scytalopus_magellanicus
Thamnophilus_nigrocinereus
Pitta_nympha
Smithornis_sharpei
Cacatua_moluccensis
Psittacus_erithacus
Aegotheles_cristatus
Phaethornis_hispidus
Rhynochetos_jubatus
Caprimulgus_indicus
Nyctibius_grandis
Heliornis_fulica
Trogon_viridis
Phodilus_badius
Tyto_alba
Strix_leptogrammica
Anhinga_rufa
Phalacrocorax_carbo
Morus_serrator
Fregata_sp
Balaeniceps_rex
Pelecanus_conspicillatus
Ciconia_ciconia
Eudocimus_ruber
Ardea_cinerea
Diomedea_chrysostoma
Larus_vegae
Eudyptula_minor
Gavia_stellata
Grus_canadensis
Cathartes_aura
Phoenicopterus_ruber
Podiceps_cristatus
Arenaria_interpres
Jacana_jacana
Haematopus_ater
Centropus_sinensis
Cuculus_poliocephalus
Crotophaga_ani
Geococcyx_californianus
Columba_livia
Geotrygon_violacea
Buteo_buteo
Pandion_haliaetus
Sagittarius_serpentarius
Falco_peregrinus
Micrastur_gilvicollis
Phaethon_lepturus
Bucorvus_leadbeateri
Dryocopus_pileatus
Upupa_epops
Alectura_lathami
Colinus_virginianus
Coturnix_japonica
Gallus_gallus
Numida_meleagris
Crax_rubra
Anas_platyrhynchos
Aythya_americana
Anser_anser
Anseranas_semipalmata
Apteryx_australis
Struthio_camelus
Casuarius_casuarius
Dromaius_novaehollandiae
Rhea_americana
Eudromia_elegans
Tinamus_guttatus
Alligator_mississippiensis
28.8351
33.8543
35.2573
20.9821
27.4221
38.6722
65.3703
42.63
32.5885
45.2004
51.2231
71.7125
62.2658
47.8683
36.6075
54.8719
37.6228
36.089
58.8425
63.5293
65.942
62.2192
67.4799
56.5442
65.4132
31.0279
57.1983
39.4031
43.0676
73.299
53.903
56.6861
39.7794
55.2618
57.0894
51.7282
58.204
47.4302
68.7715
62.6808
38.3734
52.7944
61.547
39.0324
84.2603
71.1464
63.3401
45.8621
36.5354
59.7018
30.0512
65.8821
41.6787
54.4474
43.1535
63.263
63.068
54.1571
88.7605
60.3434
34.0569
24.7551
37.4547
55.0465
70.5056
13.6484
27.7535
60.5328
225.8428
44.1196
46.3221
26.6995
53.9407
63.8162
40.111
30.0
200
150
100
50
0

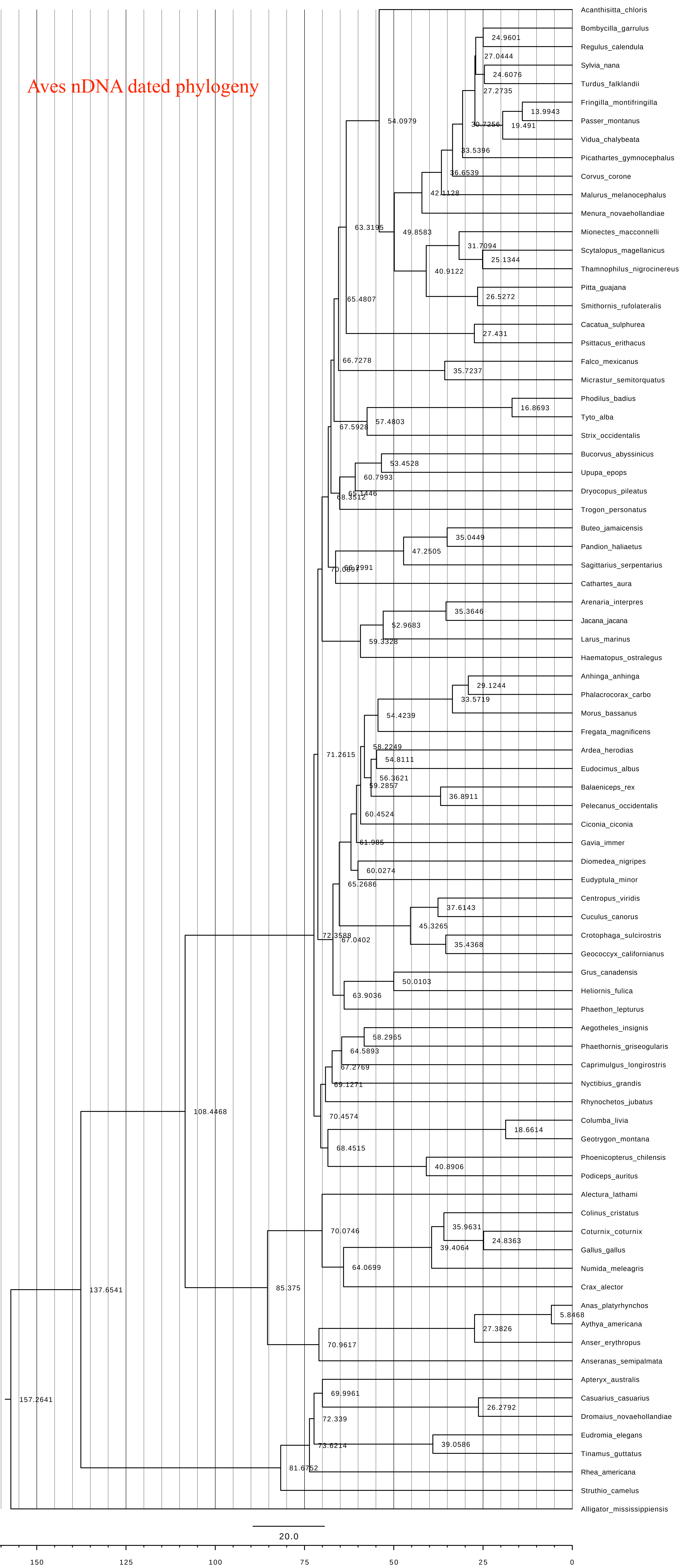
Aves nDNA dated phylogeny
Acanthisitta_chloris
Bombycilla_garrulus
Regulus_calendula
Sylvia_nana
Turdus_falklandii
Fringilla_montifringilla
Passer_montanus
Vidua_chalybeata
Picathartes_gymnocephalus
Corvus_corone
Malurus_melanocephalus
Menura_novaehollandiae
Mionectes_macconnelli
Scytalopus_magellanicus
Thamnophilus_nigrocinereus
Pitta_guajana
Smithornis_rufolateralis
Cacatua_sulphurea
Psittacus_erithacus
Falco_mexicanus
Micrastur_semitorquatus
Phodilus_badius
Tyto_alba
Strix_occidentalis
Bucorvus_abyssinicus
Upupa_epops
Dryocopus_pileatus
Trogon_personatus
Buteo_jamaicensis
Pandion_haliaetus
Sagittarius_serpentarius
Cathartes_aura
Arenaria_interpres
Jacana_jacana
Larus_marinus
Haematopus_ostralegus
Anhinga_anhinga
Phalacrocorax_carbo
Morus_bassanus
Fregata_magnificens
Ardea_herodias
Eudocimus_albus
Balaeniceps_rex
Pelecanus_occidentalis
Ciconia_ciconia
Gavia_immer
Diomedea_nigripes
Eudyptula_minor
Centropus_viridis
Cuculus_canorus
Crotophaga_sulcirostris
Geococcyx_californianus
Grus_canadensis
Heliornis_fulica
Phaethon_lepturus
Aegotheles_insignis
Phaethornis_griseogularis
Caprimulgus_longirostris
Nyctibius_grandis
Rhynochetos_jubatus
Columba_livia
Geotrygon_montana
Phoenicopterus_chilensis
Podiceps_auritus
Alectura_lathami
Colinus_cristatus
Coturnix_coturnix
Gallus_gallus
Numida_meleagris
Crax_alector
Anas_platyrhynchos
Aythya_americana
Anser_erythropus
Anseranas_semipalmata
Apteryx_australis
Casuarius_casuarius
Dromaius_novaehollandiae
Eudromia_elegans
Tinamus_guttatus
Rhea_americana
Struthio_camelus
Alligator_mississippiensis
24.9601
27.0444
24.6076
27.2735
13.9943
30.7256
19.491
54.0979
33.5396
36.6539
42.1128
63.3195
49.8583
31.7094
25.1344
40.9122
26.5272
65.4807
27.431
66.7278
35.7237
16.8693
57.4803
67.5928
53.4528
60.7993
65.1446
68.3512
35.0449
47.2505
66.2991
35.3646
52.9683
59.3328
29.1244
33.5719
54.4239
71.2615
58.2249
54.8111
56.3621
59.2857
36.8911
60.4524
61.985
60.0274
65.2686
37.6143
45.3265
72.3588
67.0402
35.4368
50.0103
63.9036
58.2965
64.5893
67.2769
69.1271
108.4468
70.4574
18.6614
68.4515
40.8906
70.0746
35.9631
24.8363
39.4064
64.0699
137.6541
85.375
5.8468
27.3826
70.9617
69.9961
26.2792
72.339
73.6214
39.0586
157.2641
81.6752
20.0
150
125
100
75
50
25
0

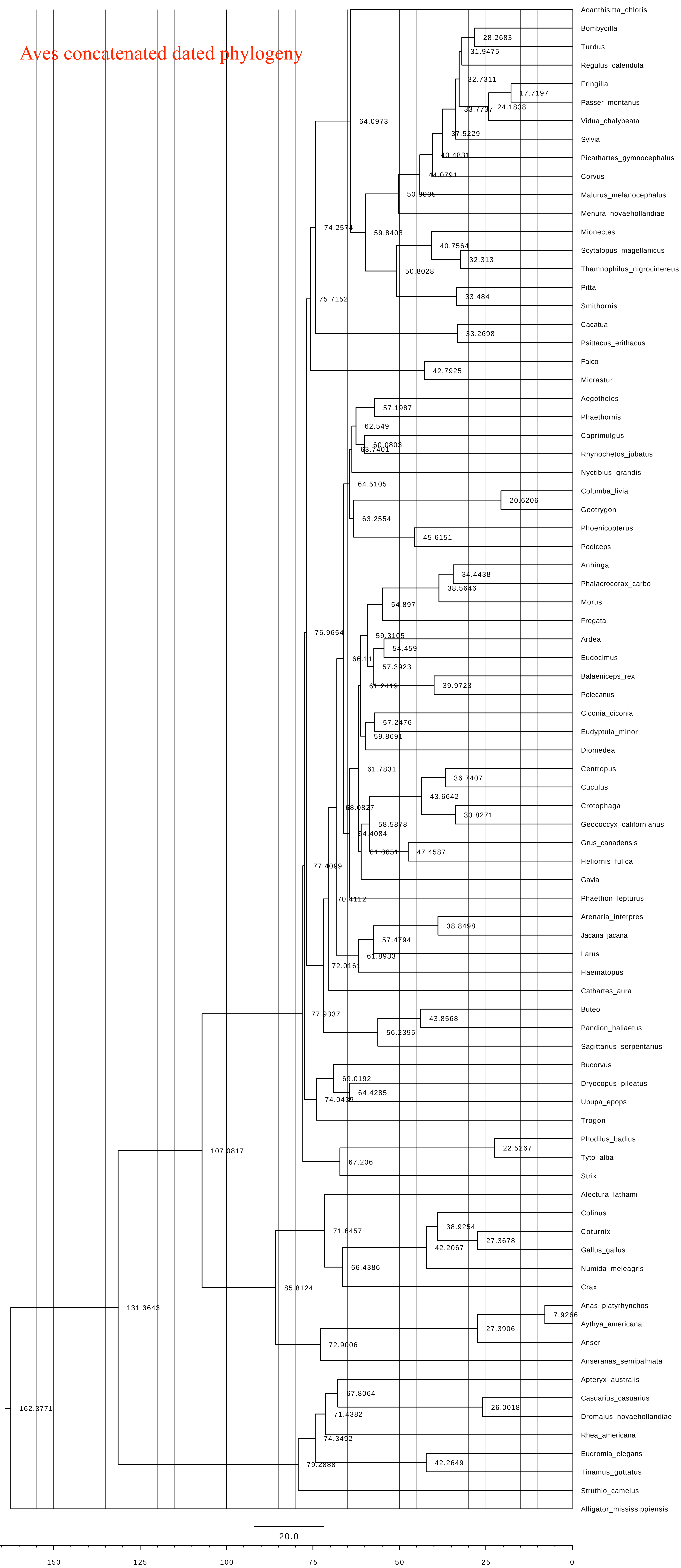

Aves concatenated dated phylogeny
Acanthisitta_chloris
Bombycilla
Turdus
Regulus_calendula
Fringilla
Passer_montanus
Vidua_chalybeata
Sylvia
Picathartes_gymnocephalus
Corvus
Malurus_melanocephalus
Menura_novaehollandiae
Mionectes
Scytalopus_magellanicus
Thamnophilus_nigrocinereus
Pitta
Smithornis
Cacatua
Psittacus_erithacus
Falco
Micrastur
Aegotheles
Phaethornis
Caprimulgus
Rhynochetos_jubatus
Nyctibius_grandis
Columba_livia
Geotrygon
Phoenicopterus
Podiceps
Anhinga
Phalacrocorax_carbo
Morus
Fregata
Ardea
Eudocimus
Balaeniceps_rex
Pelecanus
Ciconia_ciconia
Eudyptula_minor
Diomedea
Centropus
Cuculus
Crotophaga
Geococcyx_californianus
Grus_canadensis
Heliornis_fulica
Gavia
Phaethon_lepturus
Arenaria_interpres
Jacana_jacana
Larus
Haematopus
Cathartes_aura
Buteo
Pandion_haliaetus
Sagittarius_serpentarius
Bucorvus
Dryocopus_pileatus
Upupa_epops
Trogon
Phodilus_badius
Tyto_alba
Strix
Alectura_lathami
Colinus
Coturnix
Gallus_gallus
Numida_meleagris
Crax
Anas_platyrhynchos
Aythya_americana
Anser
Anseranas_semipalmata
Apteryx_australis
Casuarius_casuarius
Dromaius_novaehollandiae
Rhea_americana
Eudromia_elegans
Tinamus_guttatus
Struthio_camelus
Alligator_mississippiensis
28.2683
31.9475
32.7311
17.7197
33.7737
24.1838
37.5229
40.4831
44.0791
50.3005
64.0973
74.2574
59.8403
40.7564
32.313
50.8028
33.484
75.7152
33.2698
42.7925
57.1987
62.549
60.0803
63.7401
64.5105
20.6206
63.2554
45.6151
34.4438
38.5646
54.897
76.9654
59.3105
54.459
66.11
57.3923
61.2419
39.9723
57.2476
59.8691
61.7831
36.7407
43.6642
68.0827
33.8271
58.5878
64.4084
61.0651
47.4587
77.4099
70.4112
38.8498
57.4794
61.8933
72.0161
43.8568
77.9337
56.2395
69.0192
64.4285
74.0439
22.5267
107.0817
67.206
71.6457
38.9254
27.3678
42.2067
66.4386
85.8124
7.9266
131.3643
27.3906
72.9006
67.8064
26.0018
71.4382
162.3771
74.3492
42.2649
79.2888
20.0
150
125
100
75
50
25
0

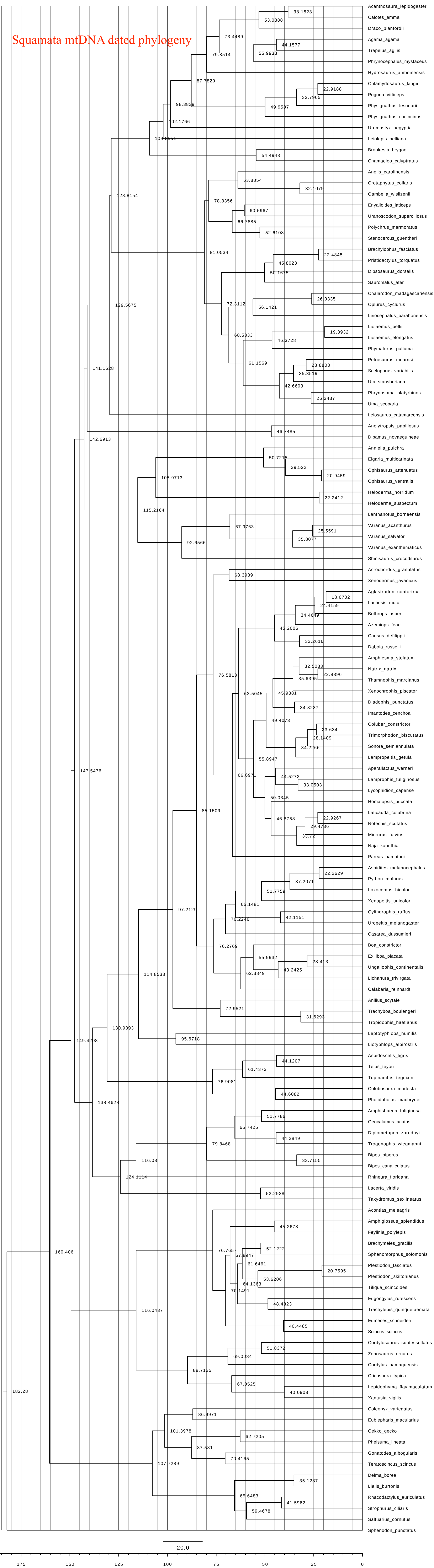
Squamata mtDNA dated phylogeny
Acanthosaura_lepidogaster
Calotes_emma
Draco_blanfordii
Agama_agama
Trapelus_agilis
Phrynocephalus_mystaceus
Hydrosaurus_amboinensis
Chlamydosaurus_kingii
Pogona_vitticeps
Physignathus_lesueurii
Physignathus_cocincinus
Uromastyx_aegyptia
Leiolepis_belliana
Brookesia_brygooi
Chamaeleo_calyptratus
Anolis_carolinensis
Crotaphytus_collaris
Gambelia_wislizenii
Enyalioides_laticeps
Uranoscodon_superciliosus
Polychrus_marmoratus
Stenocercus_guentheri
Brachylophus_fasciatus
Pristidactylus_torquatus
Dipsosaurus_dorsalis
Sauromalus_ater
Chalarodon_madagascariensis
Oplurus_cyclurus
Leiocephalus_barahonensis
Liolaemus_bellii
Liolaemus_elongatus
Phymaturus_palluma
Petrosaurus_mearnsi
Sceloporus_variabilis
Uta_stansburiana
Phrynosoma_platyrhinos
Uma_scoparia
Leiosaurus_catamarcensis
Anelytropsis_papillosus
Dibamus_novaeguineae
Anniella_pulchra
Elgaria_multicarinata
Ophisaurus_attenuatus
Ophisaurus_ventralis
Heloderma_horridum
Heloderma_suspectum
Lanthanotus_borneensis
Varanus_acanthurus
Varanus_salvator
Varanus_exanthematicus
Shinisaurus_crocodilurus
Acrochordus_granulatus
Xenodermus_javanicus
Agkistrodon_contortrix
Lachesis_muta
Bothrops_asper
Azemiops_feae
Causus_defilippii
Daboia_russelii
Amphiesma_stolatum
Natrix_natrix
Thamnophis_marcianus
Xenochrophis_piscator
Diadophis_punctatus
Imantodes_cenchoa
Coluber_constrictor
Trimorphodon_biscutatus
Sonora_semiannulata
Lampropeltis_getula
Aparallactus_werneri
Lamprophis_fuliginosus
Lycophidion_capense
Homalopsis_buccata
Laticauda_colubrina
Notechis_scutatus
Micrurus_fulvius
Naja_kaouthia
Pareas_hamptoni
Aspidites_melanocephalus
Python_molurus
Loxocemus_bicolor
Xenopeltis_unicolor
Cylindrophis_ruffus
Uropeltis_melanogaster
Casarea_dussumieri
Boa_constrictor
Exiliboa_placata
Ungaliophis_continentalis
Lichanura_trivirgata
Calabaria_reinhardtii
Anilius_scytale
Trachyboa_boulengeri
Tropidophis_haetianus
Leptotyphlops_humilis
Liotyphlops_albirostris
Aspidoscelis_tigris
Teius_teyou
Tupinambis_teguixin
Colobosaura_modesta
Pholidobolus_macbrydei
Amphisbaena_fuliginosa
Geocalamus_acutus
Diplometopon_zarudnyi
Trogonophis_wiegmanni
Bipes_biporus
Bipes_canaliculatus
Rhineura_floridana
Lacerta_viridis
Takydromus_sexlineatus
Acontias_meleagris
Amphiglossus_splendidus
Feylinia_polylepis
Brachymeles_gracilis
Sphenomorphus_solomonis
Plestiodon_fasciatus
Plestiodon_skiltonianus
Tiliqua_scincoides
Eugongylus_rufescens
Trachylepis_quinquetaeniata
Eumeces_schneideri
Scincus_scincus
Cordylosaurus_subtessellatus
Zonosaurus_ornatus
Cordylus_namaquensis
Cricosaura_typica
Lepidophyma_flavimaculatum
Xantusia_vigilis
Coleonyx_variegatus
Eublepharis_macularius
Gekko_gecko
Phelsuma_lineata
Gonatodes_albogularis
Teratoscincus_scincus
Delma_borea
Lialis_burtonis
Rhacodactylus_auriculatus
Strophurus_ciliaris
Saltuarius_cornutus
Sphenodon_punctatus
20.0
175
150
125
100
75
50
25
0

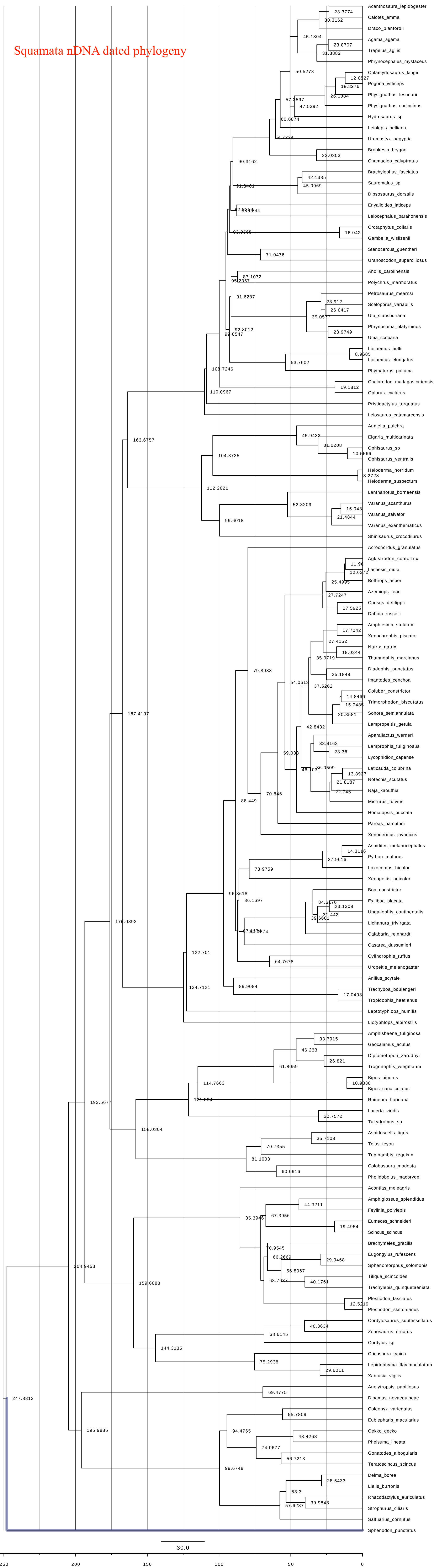

Squamata nDNA dated phylogeny
Acanthosaura_lepidogaster
23.3774
Calotes_emma
30.3162
Draco_blanfordii
45.1304
Agama_agama
23.8707
Trapelus_agilis
31.8882
Phrynocephalus_mystaceus
50.5273
Chlamydosaurus_kingii
12.0527
Pogona_vitticeps
18.8276
Physignathus_lesueurii
26.1884
57.3597
Physignathus_cocincinus
47.5392
Hydrosaurus_sp
60.6874
Leiolepis_belliana
Uromastyx_aegyptia
64.7224
Brookesia_brygooi
32.0303
Chamaeleo_calyptratus
90.3162
Brachylophus_fasciatus
42.1335
Sauromalus_sp
45.0969
91.8481
Dipsosaurus_dorsalis
Enyalioides_laticeps
92.8253
88.8244
Leiocephalus_barahonensis
Crotaphytus_collaris
93.9565
16.042
Gambelia_wislizenii
Stenocercus_guentheri
71.0476
Uranoscodon_superciliosus
Anolis_carolinensis
87.1072
95.2357
Polychrus_marmoratus
Petrosaurus_mearnsi
91.6287
28.912
Sceloporus_variabilis
26.0417
Uta_stansburiana
39.0577
Phrynosoma_platyrhinos
92.8012
23.9749
99.8547
Uma_scoparia
Liolaemus_bellii
8.9685
Liolaemus_elongatus
53.7602
Phymaturus_palluma
108.7246
Chalarodon_madagascariensis
19.1812
110.0967
Oplurus_cyclurus
Pristidactylus_torquatus
Leiosaurus_catamarcensis
163.6757
Anniella_pulchra
45.9432
Elgaria_multicarinata
31.0208
Ophisaurus_sp
10.5566
104.3735
Ophisaurus_ventralis
Heloderma_horridum
3.2728
Heloderma_suspectum
112.2621
Lanthanotus_borneensis
52.3209
Varanus_acanthurus
15.048
Varanus_salvator
21.4844
99.6018
Varanus_exanthematicus
Shinisaurus_crocodilurus
Acrochordus_granulatus
Agkistrodon_contortrix
11.96
Lachesis_muta
12.6372
Bothrops_asper
25.4995
Azemiops_feae
27.7247
Causus_defilippii
17.5925
Daboia_russelii
Amphiesma_stolatum
17.7042
Xenochrophis_piscator
27.4152
Natrix_natrix
18.0344
Thamnophis_marcianus
35.9719
Diadophis_punctatus
79.8988
25.1848
Imantodes_cenchoa
54.0613
37.5262
Coluber_constrictor
14.8466
Trimorphodon_biscutatus
15.7485
Sonora_semiannulata
20.8581
Lampropeltis_getula
42.8432
Aparallactus_werneri
33.9163
Lamprophis_fuliginosus
23.36
Lycophidion_capense
59.038
36.0509
Laticauda_colubrina
46.1031
13.8927
Notechis_scutatus
21.8187
Naja_kaouthia
22.746
Micrurus_fulvius
88.449
70.846
Homalopsis_buccata
Pareas_hamptoni
Xenodermus_javanicus
167.4197
Aspidites_melanocephalus
14.3116
Python_molurus
27.9616
Loxocemus_bicolor
78.9759
Xenopeltis_unicolor
Boa_constrictor
96.8618
86.1697
34.6176
Exiliboa_placata
23.1308
Ungaliophis_continentalis
31.442
39.6601
Lichanura_trivirgata
87.8274
82.4274
Calabaria_reinhardtii
176.0892
Casarea_dussumieri
Cylindrophis_ruffus
64.7678
122.701
Uropeltis_melanogaster
Anilius_scytale
124.7121
89.9084
Trachyboa_boulengeri
17.0403
Tropidophis_haetianus
Leptotyphlops_humilis
Liotyphlops_albirostris
Amphisbaena_fuliginosa
33.7915
Geocalamus_acutus
46.233
Diplometopon_zarudnyi
26.821
61.8059
Trogonophis_wiegmanni
114.7663
Bipes_biporus
10.9338
Bipes_canaliculatus
193.5677
121.334
Rhineura_floridana
Lacerta_viridis
30.7572
Takydromus_sp
158.0304
Aspidoscelis_tigris
35.7108
Teius_teyou
70.7355
Tupinambis_teguixin
81.1003
Colobosaura_modesta
60.0916
Pholidobolus_macbrydei
Acontias_meleagris
Amphiglossus_splendidus
44.3211
85.3946
67.3956
Feylinia_polylepis
Eumeces_schneideri
19.4954
Scincus_scincus
Brachymeles_gracilis
70.9545
Eugongylus_rufescens
66.2669
29.0468
204.9453
Sphenomorphus_solomonis
56.8067
Tiliqua_scincoides
68.7687
40.1761
159.6088
Trachylepis_quinquetaeniata
Plestiodon_fasciatus
12.5219
Plestiodon_skiltonianus
Cordylosaurus_subtessellatus
40.3634
Zonosaurus_ornatus
68.6145
Cordylus_sp
144.3135
Cricosaura_typica
75.2938
Lepidophyma_flavimaculatum
29.6011
Xantusia_vigilis
247.8812
Anelytropsis_papillosus
69.4775
Dibamus_novaeguineae
Coleonyx_variegatus
55.7809
195.9886
Eublepharis_macularius
94.4765
Gekko_gecko
48.4268
Phelsuma_lineata
74.0677
Gonatodes_albogularis
56.7213
99.6748
Teratoscincus_scincus
Delma_borea
28.5433
Lialis_burtonis
53.3
Rhacodactylus_auriculatus
57.6287
39.9848
Strophurus_ciliaris
Saltuarius_cornutus
Sphenodon_punctatus
30.0
250
200
150
100
50
0

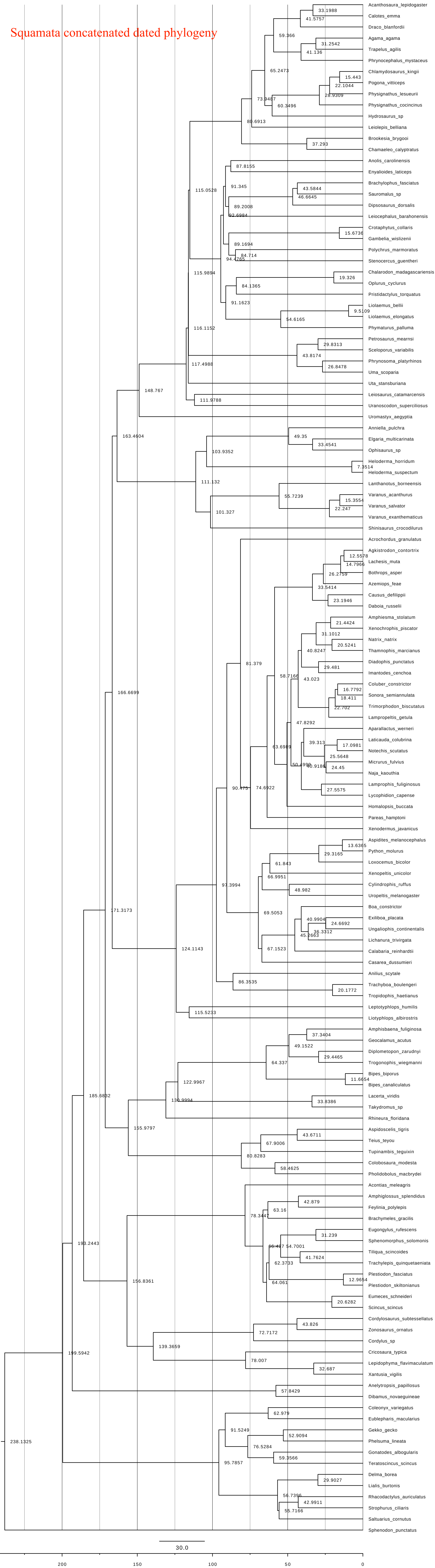
Squamata concatenated dated phylogeny
Acanthosaura_lepidogaster
Calotes_emma
Draco_blanfordii
Agama_agama
Trapelus_agilis
Phrynocephalus_mystaceus
Chlamydosaurus_kingii
Pogona_vitticeps
Physignathus_lesueurii
Physignathus_cocincinus
Hydrosaurus_sp
Leiolepis_belliana
Brookesia_brygooi
Chamaeleo_calyptratus
Anolis_carolinensis
Enyalioides_laticeps
Brachylophus_fasciatus
Sauromalus_sp
Dipsosaurus_dorsalis
Leiocephalus_barahonensis
Crotaphytus_collaris
Gambelia_wislizenii
Polychrus_marmoratus
Stenocercus_guentheri
Chalarodon_madagascariensis
Oplurus_cyclurus
Pristidactylus_torquatus
Liolaemus_bellii
Liolaemus_elongatus
Phymaturus_palluma
Petrosaurus_mearnsi
Sceloporus_variabilis
Phrynosoma_platyrhinos
Uma_scoparia
Uta_stansburiana
Leiosaurus_catamarcensis
Uranoscodon_superciliosus
Uromastyx_aegyptia
Anniella_pulchra
Elgaria_multicarinata
Ophisaurus_sp
Heloderma_horridum
Heloderma_suspectum
Lanthanotus_borneensis
Varanus_acanthurus
Varanus_salvator
Varanus_exanthematicus
Shinisaurus_crocodilurus
Acrochordus_granulatus
Agkistrodon_contortrix
Lachesis_muta
Bothrops_asper
Azemiops_feae
Causus_defilippii
Daboia_russelii
Amphiesma_stolatum
Xenochrophis_piscator
Natrix_natrix
Thamnophis_marcianus
Diadophis_punctatus
Imantodes_cenchoa
Coluber_constrictor
Sonora_semiannulata
Trimorphodon_biscutatus
Lampropeltis_getula
Aparallactus_werneri
Laticauda_colubrina
Notechis_scutatus
Micrurus_fulvius
Naja_kaouthia
Lamprophis_fuliginosus
Lycophidion_capense
Homalopsis_buccata
Pareas_hamptoni
Xenodermus_javanicus
Aspidites_melanocephalus
Python_molurus
Loxocemus_bicolor
Xenopeltis_unicolor
Cylindrophis_ruffus
Uropeltis_melanogaster
Boa_constrictor
Exiliboa_placata
Ungaliophis_continentalis
Lichanura_trivirgata
Calabaria_reinhardtii
Casarea_dussumieri
Anilius_scytale
Trachyboa_boulengeri
Tropidophis_haetianus
Leptotyphlops_humilis
Liotyphlops_albirostris
Amphisbaena_fuliginosa
Geocalamus_acutus
Diplometopon_zarudnyi
Trogonophis_wiegmanni
Bipes_biporus
Bipes_canaliculatus
Lacerta_viridis
Takydromus_sp
Rhineura_floridana
Aspidoscelis_tigris
Teius_teyou
Tupinambis_teguixin
Colobosaura_modesta
Pholidobolus_macbrydei
Acontias_meleagris
Amphiglossus_splendidus
Feylinia_polylepis
Brachymeles_gracilis
Eugongylus_rufescens
Sphenomorphus_solomonis
Tiliqua_scincoides
Trachylepis_quinquetaeniata
Plestiodon_fasciatus
Plestiodon_skiltonianus
Eumeces_schneideri
Scincus_scincus
Cordylosaurus_subtessellatus
Zonosaurus_ornatus
Cordylus_sp
Cricosaura_typica
Lepidophyma_flavimaculatum
Xantusia_vigilis
Anelytropsis_papillosus
Dibamus_novaeguineae
Coleonyx_variegatus
Eublepharis_macularius
Gekko_gecko
Phelsuma_lineata
Gonatodes_albogularis
Teratoscincus_scincus
Delma_borea
Lialis_burtonis
Rhacodactylus_auriculatus
Strophurus_ciliaris
Saltuarius_cornutus
Sphenodon_punctatus
30.0
200
150
100
50
0

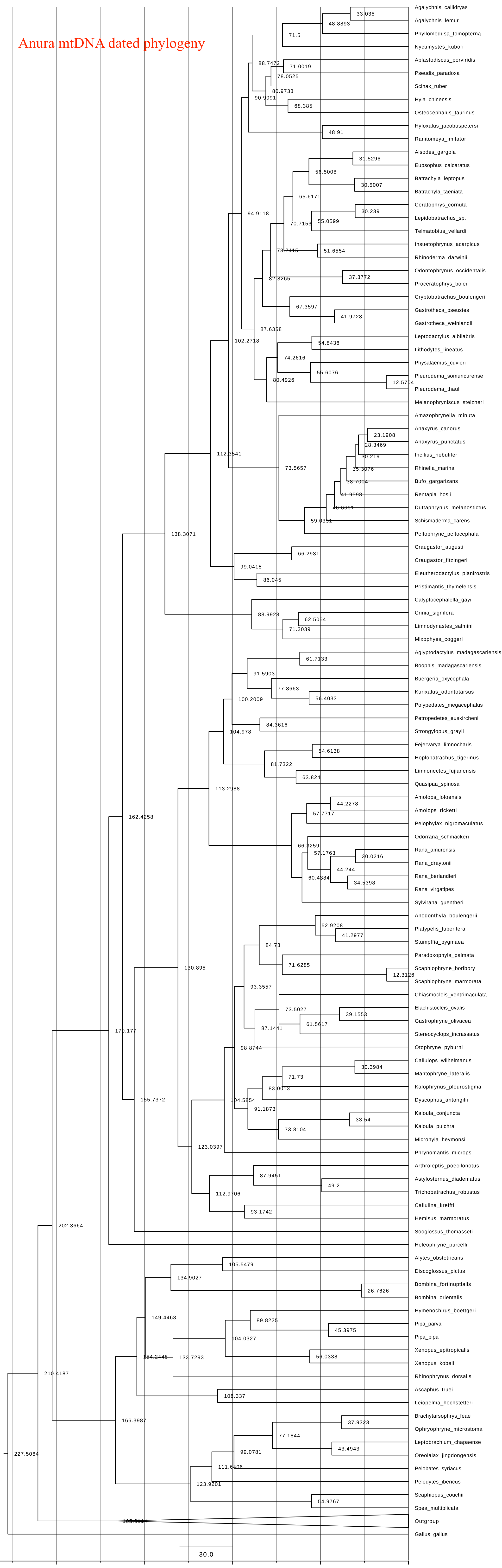

Anura mtDNA dated phylogeny
Agalychnis_callidryas
Agalychnis_lemur
Phyllomedusa_tomopterna
Nyctimystes_kubori
Aplastodiscus_perviridis
Pseudis_paradoxa
Scinax_ruber
Hyla_chinensis
Osteocephalus_taurinus
Hyloxalus_jacobuspetersi
Ranitomeya_imitator
Alsodes_gargola
Eupsophus_calcaratus
Batrachyla_leptopus
Batrachyla_taeniata
Ceratophrys_cornuta
Lepidobatrachus_sp.
Telmatobius_vellardi
Insuetophrynus_acarpicus
Rhinoderma_darwinii
Odontophrynus_occidentalis
Proceratophrys_boiei
Cryptobatrachus_boulengeri
Gastrotheca_pseustes
Gastrotheca_weinlandii
Leptodactylus_albilabris
Lithodytes_lineatus
Physalaemus_cuvieri
Pleurodema_somuncurense
Pleurodema_thaul
Melanophryniscus_stelzneri
Amazophrynella_minuta
Anaxyrus_canorus
Anaxyrus_punctatus
Incilius_nebulifer
Rhinella_marina
Bufo_gargarizans
Rentapia_hosii
Duttaphrynus_melanostictus
Schismaderma_carens
Peltophryne_peltocephala
Craugastor_augusti
Craugastor_fitzingeri
Eleutherodactylus_planirostris
Pristimantis_thymelensis
Calyptocephalella_gayi
Crinia_signifera
Limnodynastes_salmini
Mixophyes_coggeri
Aglyptodactylus_madagascariensis
Boophis_madagascariensis
Buergeria_oxycephala
Kurixalus_odontotarsus
Polypedates_megacephalus
Petropedetes_euskircheni
Strongylopus_grayii
Fejervarya_limnocharis
Hoplobatrachus_tigerinus
Limnonectes_fujianensis
Quasipaa_spinosa
Amolops_loloensis
Amolops_ricketti
Pelophylax_nigromaculatus
Odorrana_schmackeri
Rana_amurensis
Rana_draytonii
Rana_berlandieri
Rana_virgatipes
Sylvirana_guentheri
Anodonthyla_boulengerii
Platypelis_tuberifera
Stumpffia_pygmaea
Paradoxophyla_palmata
Scaphiophryne_boribory
Scaphiophryne_marmorata
Chiasmocleis_ventrimaculata
Elachistocleis_ovalis
Gastrophryne_olivacea
Stereocyclops_incrassatus
Otophryne_pyburni
Callulops_wilhelmanus
Mantophryne_lateralis
Kalophrynus_pleurostigma
Dyscophus_antongilii
Kaloula_conjuncta
Kaloula_pulchra
Microhyla_heymonsi
Phrynomantis_microps
Arthroleptis_poecilonotus
Astylosternus_diademätus
Trichobatrachus_robustus
Callulina_kreffti
Hemisus_marmoratus
Sooglossus_thomasseti
Heleophryne_purcelli
Alytes_obstetricans
Discoglossus_pictus
Bombina_fortinuptialis
Bombina_orientalis
Hymenochirus_boettgeri
Pipa_parva
Pipa_pipa
Xenopus_epitropicalis
Xenopus_kobeli
Rhinophrynus_dorsalis
Ascaphus_truei
Leiopelma_hochstetteri
Brachytarsophrys_feae
Ophryophryne_microstoma
Leptobrachium_chapaense
Oreolalax_jingdongensis
Pelobates_syriacus
Pelodytes_ibericus
Scaphiopus_couchii
Spea_multiplicata
Outgroup
Gallus_gallus
33.035
48.8893
71.5
88.7472
71.0019
78.0525
80.9733
90.9091
68.385
48.91
31.5296
56.5008
30.5007
65.6171
94.9118
30.239
70.7153
55.0599
78.2415
51.6554
82.8265
37.3772
67.3597
41.9728
87.6358
102.2718
54.8436
74.2616
55.6076
12.5704
80.4926
112.3541
73.5657
23.1908
28.3469
30.219
35.3076
38.7004
41.9598
46.6661
59.0351
138.3071
66.2931
99.0415
86.045
88.9928
62.5054
71.3039
61.7133
91.5903
77.8663
100.2009
56.4033
104.978
84.3616
54.6138
81.7322
63.824
113.2988
44.2278
57.7717
162.4258
66.3259
57.1763
30.0216
44.244
60.4384
34.5398
52.9208
41.2977
84.73
71.6285
130.895
12.3126
93.3557
73.5027
39.1553
61.5617
87.1441
170.177
98.8744
30.3984
71.73
83.0013
104.5854
155.7372
91.1873
33.54
73.8104
123.0397
87.9451
49.2
112.9706
93.1742
202.3664
105.5479
134.9027
26.7626
149.4463
89.8225
45.3975
104.0327
154.2448
133.7293
56.0338
210.4187
108.337
166.3987
37.9323
77.1844
43.4943
99.0781
227.5064
111.6406
123.9201
54.9767
165.9114
30.0
200
150
100
50
0

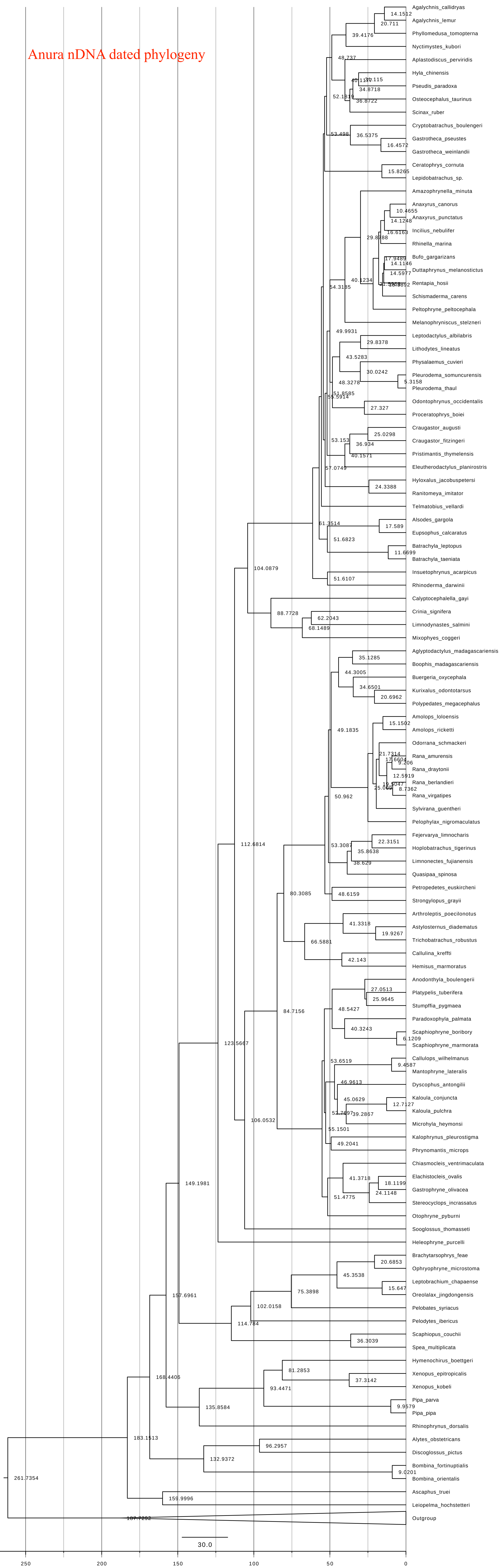

Anura nDNA dated phylogeny
Agalychnis_callidryas
Agalychnis_lemur
Phyllomedusa_tomopterna
Nyctimystes_kubori
Aplastodiscus_perviridis
Hyla_chinensis
Pseudis_paradoxa
Osteocephalus_taurinus
Scinax_ruber
Cryptobatrachus_boulengeri
Gastrotheca_pseustes
Gastrotheca_weinlandii
Ceratophrys_cornuta
Lepidobatrachus_sp.
Amazophrynella_minuta
Anaxyrus_canorus
Anaxyrus_punctatus
Incilius_nebulifer
Rhinella_marina
Bufo_gargarizans
Duttaphrynus_melanostictus
Rentapia_hosii
Schismaderma_carens
Peltophryne_peltocephala
Melanophryniscus_stelzneri
Leptodactylus_albilabris
Lithodytes_lineatus
Physalaemus_cuvieri
Pleurodema_somuncurensis
Pleurodema_thaul
Odontophrynus_occidentalis
Proceratophrys_boiei
Craugastor_augusti
Craugastor_fitzingeri
Pristimantis_thymelensis
Eleutherodactylus_planirostris
Hyloxalus_jacobuspetersi
Ranitomeya_imitator
Telmatobius_vellardi
Alsodes_gargola
Eupsophus_calcaratus
Batrachyla_leptopus
Batrachyla_taeniata
Insuetophrynus_acarpicus
Rhinoderma_darwinii
Calyptocephalella_gayi
Crinia_signifera
Limnodynastes_salmini
Mixophyes_coggeri
Aglyptodactylus_madagascariensis
Boophis_madagascariensis
Buergeria_oxycephala
Kurixalus_odontotarsus
Polypedates_megacephalus
Amolops_loloensis
Amolops_ricketti
Odorrana_schmackeri
Rana_amurensis
Rana_draytonii
Rana_berlandieri
Rana_virgatipes
Sylvirana_guentheri
Pelophylax_nigromaculatus
Fejervarya_limnocharis
Hoplobatrachus_tigerinus
Limnonectes_fujianensis
Quasipaa_spinosa
Petropedetes_euskircheni
Strongylopus_grayii
Arthroleptis_poecilonotus
Astylosternus_diadematus
Trichobatrachus_robustus
Callulina_kreffti
Hemisus_marmoratus
Anodonthyla_boulengerii
Platypelis_tuberifera
Stumpffia_pygmaea
Paradoxophyla_palmata
Scaphiophryne_boribory
Scaphiophryne_marmorata
Callulops_wilhelmanus
Mantophryne_lateralis
Dyscophus_antongilii
Kaloula_conjuncta
Kaloula_pulchra
Microhyla_heymonsi
Kalophrynus_pleurostigma
Phrynomantis_microps
Chiasmocleis_ventrimaculata
Elachistocleis_ovalis
Gastrophryne_olivacea
Stereocyclops_incrassatus
Otophryne_pyburni
Sooglossus_thomasseti
Heleophryne_purcelli
Brachytarsophrys_feae
Ophryophryne_microstoma
Leptobrachium_chapaense
Oreolalax_jingdongensis
Pelobates_syriacus
Pelodytes_ibericus
Scaphiopus_couchii
Spea_multiplicata
Hymenochirus_boettgeri
Xenopus_epitropicalis
Xenopus_kobeli
Pipa_parva
Pipa_pipa
Rhinophrynus_dorsalis
Alytes_obstetricans
Discoglossus_pictus
Bombina_fortinuptialis
Bombina_orientalis
Ascaphus_truei
Leiopelma_hochstetteri
Outgroup
30.0
250
200
150
100
50
0

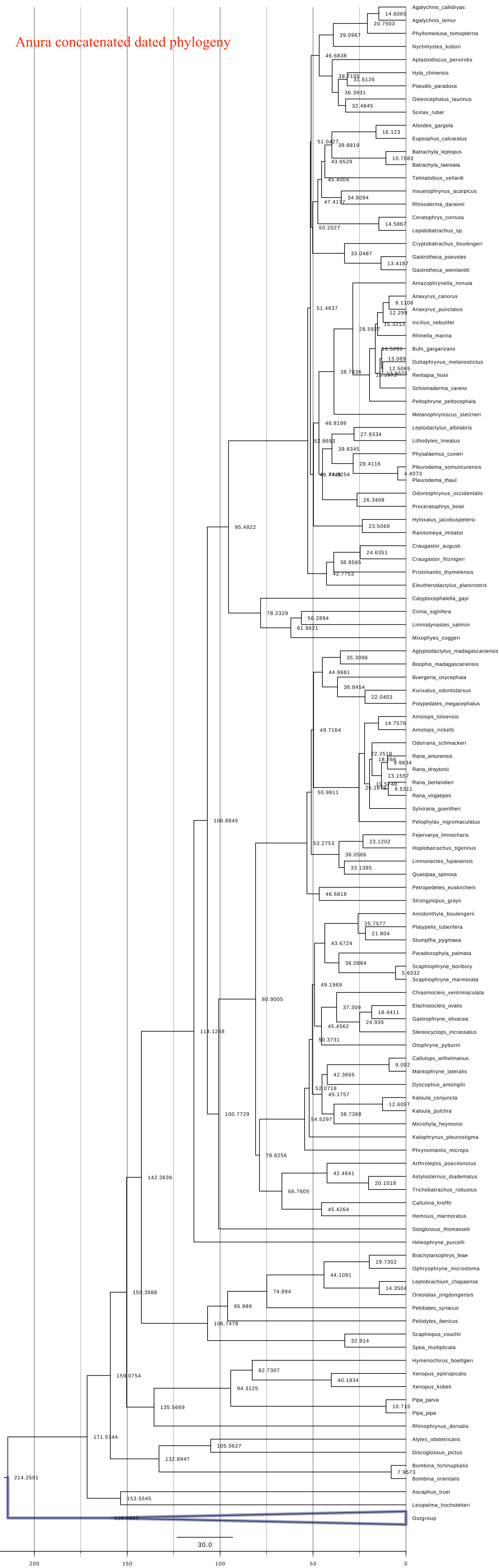

Anura concatenated dated phylogeny
Agalychnis_callidryas
Agalychnis_lemur
Phyllomedusa_tomopterna
Nyctimystes_kubori
Aplastodiscus_perviridis
Hyla_chinensis
Pseudis_paradoxa
Osteocephalus_taurinus
Scinax_ruber
Alsodes_gargola
Eupsophus_calcaratus
Batrachyla_leptopus
Batrachyla_taeniata
Telmatobius_vellardi
Insuetophrynus_acarpicus
Rhinoderma_darwinii
Ceratophrys_cornuta
Lepidobatrachus_sp.
Cryptobatrachus_boulengeri
Gastrotheca_pseustes
Gastrotheca_weinlandii
Amazophrynella_minuta
Anaxyrus_canorus
Anaxyrus_punctatus
Incilius_nebulifer
Rhinella_marina
Bufo_gargarizans
Duttaphrynus_melanostictus
Rentapia_hosii
Schismaderma_carens
Peltophryne_peltocephala
Melanophryniscus_stelzneri
Leptodactylus_albilabris
Lithodytes_lineatus
Physalaemus_cuvieri
Pleurodema_somuncurensis
Pleurodema_thaul
Odontophrynus_occidentalis
Proceratophrys_boiei
Hyloxalus_jacobuspetersi
Ranitomeya_imitator
Craugastor_augusti
Craugastor_fitzingeri
Pristimantis_thymelensis
Eleutherodactylus_planirostris
Calyptocephalella_gayi
Crinia_signifera
Limnodynastes_salmini
Mixophyes_coggeri
Aglyptodactylus_madagascariensis
Boophis_madagascariensis
Buergeria_oxycephala
Kurixalus_odontotarsus
Polypedates_megacephalus
Amolops_loloensis
Amolops_ricketti
Odorrana_schmackeri
Rana_amurensis
Rana_draytonii
Rana_berlandieri
Rana_virgatipes
Sylvirana_guentheri
Pelophylax_nigromaculatus
Fejervarya_limnocharis
Hoplobatrachus_tigerinus
Limnonectes_fujianensis
Quasipaa_spinosa
Petropedetes_euskircheni
Strongylopus_grayii
Anodonthyla_boulengerii
Platypelis_tuberifera
Stumpffia_pygmaea
Paradoxophyla_palmata
Scaphiophryne_boribory
Scaphiophryne_marmorata
Chiasmocleis_ventrimaculata
Elachistocleis_ovalis
Gastrophryne_olivacea
Stereocyclops_incrassatus
Otophryne_pyburni
Callulops_wilhelmanus
Mantophryne_lateralis
Dyscophus_antongilii
Kaloula_conjuncta
Kaloula_pulchra
Microhyla_heymonsi
Kalophrynus_pleurostigma
Phrynomantis_microps
Arthroleptis_poecilonotus
Astylosternus_diadematus
Trichobatrachus_robustus
Callulina_kreffti
Hemisus_marmoratus
Sooglossus_thomasseti
Heleophryne_purcelli
Brachytarsophrys_feae
Ophryophryne_microstoma
Leptobrachium_chapaense
Oreolalax_jingdongensis
Pelobates_syriacus
Pelodytes_ibericus
Scaphiopus_couchii
Spea_multiplicata
Hymenochirus_boettgeri
Xenopus_epitropicalis
Xenopus_kobeli
Pipa_parva
Pipa_pipa
Rhinophrynus_dorsalis
Alytes_obstetricans
Discoglossus_pictus
Bombina_fortinuptialis
Bombina_orientalis
Ascaphus_truei
Leiopelma_hochstetteri
Outgroup
30.0
200
150
100
50
0
